# Synovium-Restricted Armored PD-1-Targeted CAR-T Cells Reprogram Immunity and Resolve Experimental Arthritis

**DOI:** 10.64898/2026.07.09.737520

**Authors:** Chamutal Gur, Lior Ravkaie, Reut Sharet-Eshed, Rotem Shalita, Roberto Avellino, Einat Rauchbach, Ken Xie, Eyal David, Gal Yagel, Mor Zada, Maya Ben Yehuda, Kfir Mazuz, Michelle Nathalie Von Locquenghien, Hagit Peleg, Yaakov Naparstek, Karine Atlan, Shlomit Kfir-Erenfeld, Yuri Kuznetsov, Reut Tzemach, Merav Lidar, Alexandra Balbir-Gurman, Truong San Phan, Kiara Freitag, Ido Amit

## Abstract

Despite major therapeutic advances, a substantial fraction of patients with autoimmune disease remains refractory to treatment. While B cell-targeted CAR-T therapies have shown considerable efficacy, the central contribution of pathogenic T cells to rheumatoid arthritis (RA) suggests that complementary T cell-directed strategies may enable deeper disease control. Using single-cell multi-omics of human RA and experimental models, *PDCD1* was identified as a selective marker of synovial disease-associated T cells. We developed PD-1-directed CAR-T cells that potently eliminate these cells *in vitro* and *in vivo*, leading to marked attenuation of synovitis in RA models. To limit off-target activity, we engineered NR4A2-driven CAR-responsive biosensors to restrict CAR activity to inflamed synovium. To couple anti-PD-1 CAR-mediated cytotoxicity with microenvironmental modulation, we further engineered these CAR-T cells to secrete soluble TNF receptor II (sTNFRii), counteracting baseline inflammation and CAR-induced IFN response and promoting a tissue-reparative myeloid state. PD-1-targeted CAR-T therapy thus represents a promising, specific, and safe strategy for autoimmune diseases involving disease-associated T cells.

## Introduction

Autoimmune diseases affect millions worldwide and remain a major cause of chronic morbidity despite advances in immunomodulatory therapies. Across many autoimmune conditions, 10-30% of patients remain refractory to standard treatments or experience recurrent relapses, placing them at risk of progressive organ damage, disability, and increased mortality^1–7^. Moreover, many current therapies require prolonged or repeated administration, as they primarily suppress inflammation rather than reliably restoring durable immune tolerance, and may be associated with adverse effects.

Chimeric antigen receptor (CAR)-T cell therapy, originally developed for hematologic malignancies, is emerging as a promising therapeutic approach for autoimmune diseases^8–16^. Clinical success has been most striking in disorders with prominent B cell involvement, such as systemic lupus erythematosus, dermatomyositis, and myasthenia gravis, where CD19- or BCMA-directed CAR-T cells can induce profound B cell depletion, immune resetting, and durable remission in severe treatment-refractory cases^8,11,12,17,18^.

These advances suggest that CAR-T therapy may represent a transformative modality in autoimmunity. However, they also point to a potential conceptual limitation: current CAR-T approaches for autoimmune diseases are primarily directed against B cells^8,12,17^, whereas many autoimmune pathologies are not exclusively B cell-mediated. This consideration may be particularly relevant in diseases characterized by extensive B and T cell crosstalk, such as rheumatoid arthritis (RA)^19–21^. Although B cells contribute to disease pathogenesis through autoantibody production, antigen presentation, and cytokine secretion^21–23^, disease-associated T-cell populations are central to the initiation, propagation, and persistence of tissue inflammation ^20,21,24–26^. T cells not only amplify local inflammatory circuits through myeloid attraction and activation, but also sustain B cell activation, organize ectopic lymphoid structures, and shape pathogenic interactions between myeloid and stromal cells ^20,21,24,25^. Thus, in some disease settings, targeting B cells alone may leave intact the cellular programs that maintain chronic inflammation, tissue residency, and immunological memory. In this context, T cell-directed CAR-T therapy may represent an opportunity to target diseases in which pathogenic T cells act as core organizers of the inflamed tissue niche.

Additional challenges remain for CAR-T therapy in autoimmune diseases, including limited specificity for pathogenic immune populations and unintended depletion of non-pathogenic cells ^27,28^. Importantly, relapses after B cell-directed CAR-T therapy in autoimmune diseases^29–31^ indicate that durable immune resetting is not always achieved and suggest that other disease-driving mechanisms may persist. More broadly, the factors governing CAR-T cell trafficking, tissue infiltration, retention within inflamed sites, persistence, and phenotypic stability remain incompletely understood, particularly in non-malignant inflammatory settings. These issues are of high priority when targeting disease-associated, tissue-resident T cell populations that reside within complex pathogenic tissue microenvironments.

RA is often considered a prototypic example of such a disease. Although early reports suggest that CD19-targeted CAR-T therapy may be effective in RA, the available evidence remains limited, cytokine release syndrome (CRS) and immune effector cell-associated neurotoxicity syndrome (ICANS) have been reported, including isolated higher-grade events, and key questions regarding durability and relapse have yet to be fully addressed^32–34^. At the same time, a substantial body of genetic, cellular, and tissue-level evidence supports a central role for T cells in RA pathogenesis. Susceptibility loci such as *HLA-DRB1*, *PTPN22*, and *STAT4* point to dysregulated T cell activation as a major axis of disease risk^35,36^, while synovial tissue harbors clonally expanded T cell populations with restricted T cell receptor repertoires^37–39^. These observations suggest that antigen-experienced T cells are not passive bystanders, but active drivers of local RA biology.

Recent multi-omics studies integrating single-cell and spatial profiling have further refined this view by characterizing disease-associated T cell states in inflamed RA synovium. These include clonally expanded inflammatory CD4⁺ T cells producing mediators such as IL-17, IFN-γ, and GM-CSF^21,24,37,40^, as well as peripheral helper T (Tph) cells, which provide potent B cell help, and promote local autoantibody production and ectopic lymphoid structure formation^26,38,40,41^.

Notably, accessible chromatin regions in Tph cells are enriched for RA risk variants^42^. In addition, disease-associated T cell states include cytotoxic or tissue-adapted populations, including GZMK⁺ effector-memory CD8⁺ T cells linked to chronic antigen exposure^21,43^. Together, these findings support a model in which multiple disease-associated T cell states cooperate to sustain synovial inflammation and reinforce pathogenic immune circuits. Despite this increasingly detailed cellular map, the functional contribution of individual T cell subsets to RA persistence remains incompletely resolved. This uncertainty has limited the development of precision strategies to selectively eliminate the T cell populations that sustain disease while sparing broader protective immunity. Collectively, these findings indicate that an effective cellular therapy for RA requires more than suppressing inflammation: it necessitates elimination of disease-associated synovial T cells, disruption of pathogenic T-B cell crosstalk, and reprogramming of the inflamed tissue environment toward homeostasis. These considerations provide a strong rationale for the development of CAR-T approaches targeting disease-associated T cells.

In this study, integrative multi-omics analysis of human RA samples identified disease-associated synovial T cell populations, including Tph and effector-memory CD8⁺ T cells, and prioritized *PDCD1* as a selective surface marker across these populations. We developed and evaluated a novel anti-PD-1 CAR-T strategy in complementary arthritis models, demonstrating that targeting PD-1⁺ disease-associated T cells significantly mitigates disease, disrupts pathogenic T-B interactions, and reshapes the synovial immune environment. To further improve selectivity and therapeutic function, we incorporated synovium-responsive regulatory circuits and sTNFRii cargo delivery, enabling local reinforcement of anti-inflammatory tissue remodeling. Together, our findings establish T cell-directed CAR-T therapy as a promising strategy for autoimmune diseases such as RA, in which disease-associated T cells are central drivers of chronic inflammation and tissue pathology.

## Results

### PD-1 is selectively expressed by disease-associated synovial T cells in human and murine RA

To comprehensively characterize disease-associated T cells as potential targets for CAR-T cell therapy in RA, we collected, curated, and generated scRNA-seq data from healthy and diseased human and murine synovial tissues, as well as from human peripheral blood mononuclear cells (PBMCs) and murine splenic tissues (Fig. 1a). Our human atlas integrated data from 89 RA synovial and PBMCs samples across multiple studies and disease states, including treatment-naive patients and those with an inadequate response to methotrexate or TNF inhibition, together with 9 osteoarthritis (OA) synovial samples as non-inflammatory controls and 18 PBMCs samples from healthy individuals^21,44^. After quality control, the dataset comprised 95,479 high-quality T cells, including 88,388 cells from RA samples, 4,323 cells from control OA synovium, and 2,768 cells from healthy individuals (Fig. 1a). Cells were integrated using scVI^45^ and clustered in the latent space with the Leiden algorithm (Fig. 1b, c and Supplementary Fig. 1a-c). Cell identities were annotated based on the normalized expression of canonical marker genes (Supplementary Fig. 1d-e).

**Figure 1.**
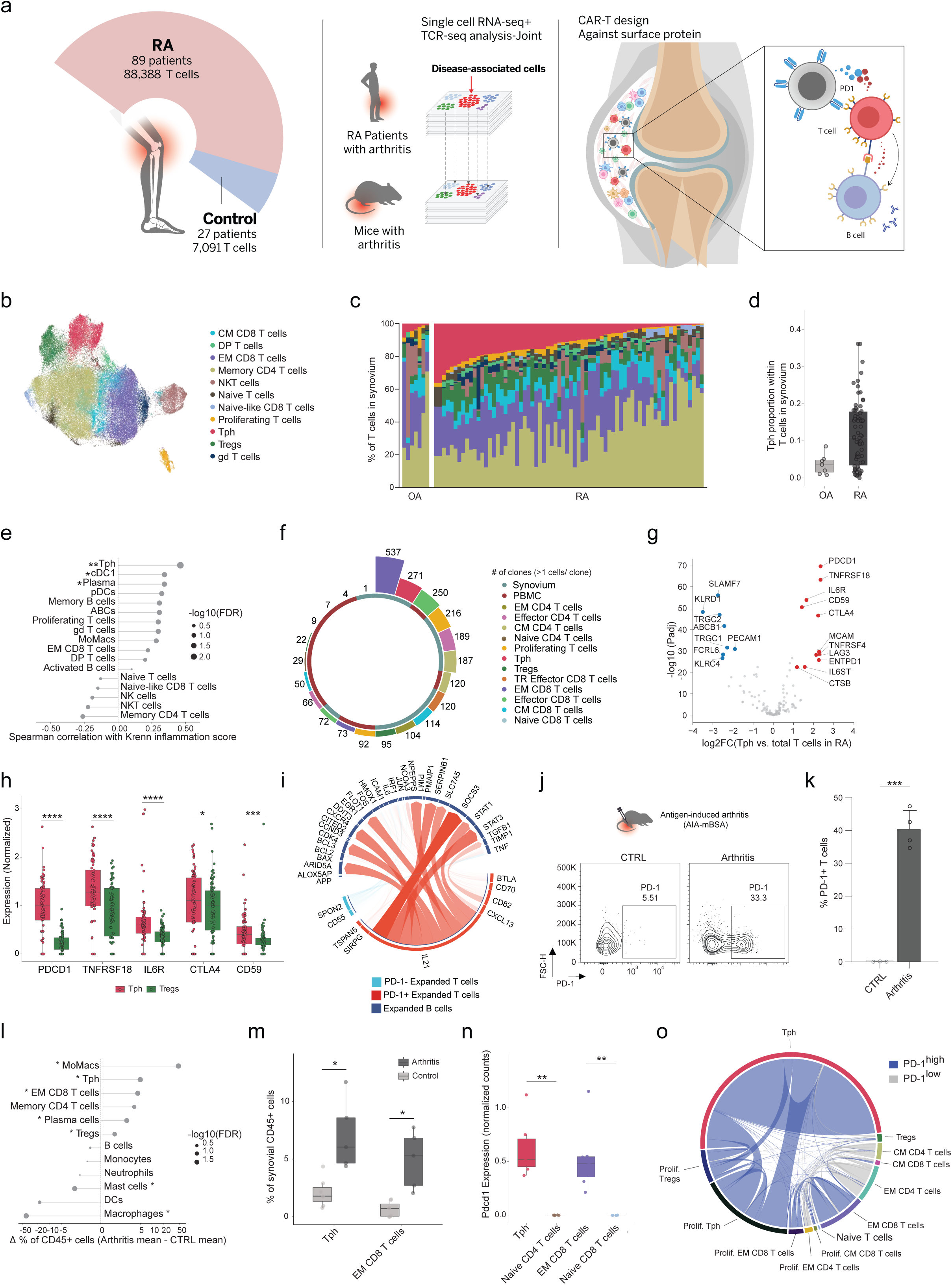
*PDCD1* is highly expressed in disease-associated synovial T cells in both human and murine samples. (a) Schematic overview of data analysis of synovial tissue from human and murine models and CAR design. (b) Integrated T cells UMAP, with graph-based clustering of 95,479 T cells from the synovium and PBMC of 116 patients (89 RA, 9 OA, 18 healthy). (c) Bar graphs depicting the percentages of synovial T cell subsets per individual grouped by disease. Patients with less than 50 T cells were excluded. (d) Boxplot showing the percentage of synovial Tph out of total synovial T cells across diseases. Each point represents a patient. (e) Spearman’s correlation testing between Krenn inflammation score and percentages of synovial T cell subpopulation in RA patients, for subpopulations with a correlation higher than 0.1. Dot size represents the statistical significance of the correlation. (** P < 0.01), (* P < 0.05). (f) Circular bar plot of the number of cells with TCR clone size higher than 1, per T subpopulation, in 12 synovial samples and 10 PBMC samples of RA patients. (g) Volcano plot of DEGs in Tph compared to other synovial T cells in RA patients. Only genes encoding plasma-membrane proteins by GO cellular-component annotations were included. Top genes with adjusted P < 10^−20^ and |log₂FC| > 1 were highlighted. (h) Boxplot showing normalized expression of top 5 genes from (g) in Tph compared to Tregs. Each point represents an RA patient. (**** P < 0.0001), (*** P < 0.001), (* P < 0.05). (i) Chord diagram of 40 strongest ligand-receptor interaction network between synovial expanded T cells as ligand-expressing senders, divided by PDCD1 raw counts, and synovial expanded B cells as receptor-expressing receivers. Transparency reflects interaction strength. (j) Representative flow cytometry analysis of PD-1 expression in T cells from healthy and inflamed synovial tissue. T cells were gated from the lymphocyte population (CD45.2⁺CD90.2⁺). (k) Summary of flow cytometry analysis of PD-1 staining from multiple mice (n=5). (*** P < 0.001). (l) Difference in average cell-type percentage of CD45^+^ immune cells in the synovium of arthritis mice compared to controls. Dot size represents the statistical significance of the difference. (* P < 0.05). (m) Boxplot showing the percentage of synovial Tph and EM CD8^+^ T cells of total synovial T cells in arthritis mice (n=5) and controls (n=6). Each point represents a mouse. (* P < 0.05). (n) Boxplot showing normalized expression of Pdcd1 in selected synovial T cell subsets. Each point represents an arthritis mouse (n=6). (** P < 0.01). (o) Chord diagram showing the interconnectivity between T cells sharing the same CDR3 (paired TCRα, and TCRβ chains) amino acid sequences (clone) within synovial T-cell PD-1^high^ and PD-1^low^ clusters (n=22 inflamed synovial tissues). Number of clones was normalized by total cells in each sample.

T cell annotations were integrated with those of other immune compartments, resulting in an atlas comprising a broad range of immune cell types, including macrophages, monocytes, dendritic cells (DCs), plasmacytoid dendritic cells (pDCs), B cells, plasma cells, NK cells, mast cells, and 11 T cells subsets, including: central memory CD8 (CM), effector memory CD8 (EM), memory CD4, NK-like CD8, NKT, naive CD4 and CD8, proliferating, T peripheral helper (Tph), regulatory T cells (Tregs), and gamma-delta (gd) T cells (Fig. 1b, c, and Supplementary Fig. 1a-e). Cell type abundance analysis identified marked differences in immune composition between RA and OA synovial tissue, along with inter-patient variability (Fig. 1c, Supplementary Fig. 1b).

RA synovial samples were predominantly enriched for T cells, alongside B cells, NK cells, and monocyte-derived macrophages (MoMacs), whereas OA samples showed enrichment of tissue-resident macrophages and reduced lymphocyte populations (Supplementary Fig. 1b). In contrast, the cellular composition of PBMCs in RA patients did not reveal any significant differences compared to healthy controls, emphasizing the relevance of synovial T cells in RA (Supplementary Fig. 1c). To identify T cell populations with the highest therapeutic potential, we focused on subsets enriched in synovial tissue that exhibit clonal expansion, inflammatory transcriptional programs, features consistent with disease pathology, and strong cross-compartment interactions. Notably, RA synovial samples exhibited a significantly higher proportion of Tph cells, which strongly correlated with the degree of synovial tissue inflammation, as assessed by the Krenn inflammation score^46^ (Fig. 1d-e), with a concurrent increase in Treg cells and a trend toward higher frequencies of *GZMK*+ EM CD8⁺ T cells (Supplementary Fig. 1e-f). Building on cellular enrichment patterns and their association with RA pathology, we integrated published paired scRNA-seq, TCR, and BCR-seq datasets of both synovial tissues and PBMCs of RA patients^37^ (Fig. 1f, Supplementary Fig. 1g-h, Supplementary Fig. 2a-g). This analysis identified Tph cells as a dominant, highly activated CD4⁺ effector memory subset in synovial tissue, exhibiting marked clonal expansion (Fig. 1f, Supplementary Fig. 2a-b). Within the CD8⁺ synovial compartment, the EM subset constituted the most clonally expanded population, consistent with ongoing antigen-specific stimulation (Fig. 1f, Supplementary Fig. 2a-b). Furthermore, Tph and EM CD8⁺ T cells were significantly enriched in synovial tissue relative to PBMCs from patients with RA, indicating tissue-specific expansion (Supplementary Fig. 2h).

Within the synovial B cell compartment, activated B cells and plasma cells were the most enriched and clonally expanded populations (Supplementary Fig. 2c-d, Supplementary Fig. 2f-g). Notably, scRNA-seq analyses revealed strong correlations between Tph cells and various B cell subsets, particularly age-associated B cells (ABCs) and plasma cells, supporting a functional link between these populations within the inflamed synovial microenvironment (Supplementary Fig. 1i-k). Collectively, these findings position Tph, together with the EM CD8⁺ T cell subset, as key mediators of disease and compelling targets for selective CAR-T therapy in RA. We leveraged our dataset to identify a cell-surface target on disease-associated T cells suitable for T cell-directed therapy. Selection criteria were: (1) membrane localization; (2) selective or enriched expression in pathogenic populations, with minimal expression in regulatory and other non-pathogenic subsets; (3) high and consistent expression across patients within affected tissue; and (4) functional relevance, defined by association with key disease-driving processes. We identified several candidate genes, with *PDCD1* showing the highest and most specific membrane expression on synovial Tph cells (Fig. 1g). *PDCD1* was also expressed at lower levels in EM CD8^+^ and proliferating T cells (Supplementary Fig. 2e). Other genes highly expressed in Tph cells (e.g., *IL*6*R*, *TNFRSF18*, *CD59* and *CTLA4*) were less specific, as they were also expressed in Tregs (Fig. 1h).

We next performed cell-cell interaction analysis using synovial expanded versus non-expanded B cells as receiver populations and PD-1^+^ versus PD-1^-^ synovial expanded T cells as sender populations. The top 40 predicted interactions were predominantly enriched for canonical Tph and activated B cell markers (Fig. 1i), further supporting a central role for PD-1^+^ T cells in disease pathogenesis. Overall, synovial-resident Tph along with EM CD8⁺ T cells were identified as clonally expanded and chronically activated contributors to human RA, consistent with ongoing antigen-driven responses. Their distinct surface expression of *PDCD1* highlights PD-1 as a potential disease specific target for CAR-T therapy in humans.

To assess the therapeutic potential of our T cell target in RA and define T cell dynamics following therapy, we performed multi-omics analyses of several murine RA models, including the mBSA-induced RA model. This antigen-driven, T cell-mediated model shows pronounced expansion of PD1⁺ T cells following arthritis, particularly during the re-challenge phase (“relapse”, Fig. 1j-k, Supplementary Fig. 3a-c, Supplementary Fig. 3e-f), mirroring human RA. The model provides a synchronized, reproducible disease course for controlled analysis of T cell dynamics. It recapitulates key features of human RA, including synovial hyperplasia, leukocyte infiltration, cartilage degradation, and bone erosion^47,48^, supporting its use in studying T cell-driven inflammation. Following arthritis induction during the re-challenge phase, synovial tissues were isolated from both knee and ankle joints, dissociated into single-cell suspensions and fluorescence-activated index cell sorting (FACS) was performed for MARS-seq profiling^49,50^. Cells were analyzed using PCA and clustered with the Leiden algorithm (Supplementary Fig. 3a-b), followed by annotation based on the normalized expression of canonical marker genes (Supplementary Fig. 3d). Consistent with human data, synovial tissue from inflamed joints in the mBSA arthritis model showed marked enrichment of T cells and MoMacs, whereas non-inflamed synovial samples were enriched in tissue-resident macrophages (Supplementary Fig. 3e-f). Within the T cell compartment, Tph and EM CD8⁺ T cells exhibited high *Pdcd1* expression, and together with Treg subsets (expressing lower levels of *Pdcd1*), were preferentially abundant and transcriptionally distinct from those in non-arthritic synovial samples, or synovial naïve T cells (Fig. 1l-n, Supplementary Fig. 3f-g).

To further resolve disease-associated T cell populations in this model, we combined TCR-seq with scRNA-seq of synovial T cells from 22 inflamed synovial samples collected following arthritis re-challenge. Synovial T cells were sorted into PD-1^high^ and PD-1^low^ populations (Supplementary Fig. 3h). The PD-1^high^ gate was predominantly composed of approximately 70% Tph cells, including a substantial fraction of proliferating Tph cells, and EM CD8⁺ T cells, whereas the PD-1^low^ gate was composed primarily of EM CD4⁺ T cells, memory CD4⁺ T cells, and Tregs (Fig. 1o, Supplementary Fig. 3h-j). TCR-seq analysis identified dominant clonotypes enriched in the PD-1^high^ population, arising predominantly from PD-1^high^ Tph and proliferating Tph cells, with a smaller contribution from EM CD8⁺ T cells and proliferating Tregs. In contrast, the PD-1^low^ population displayed a diverse TCR repertoire with no dominant clones (Fig. 1o), highlighting the PD-1^high^ compartment as the principal site of pathogenic clonal expansion and supporting its central pathogenic role in RA. Our integrative multi-omic approach demonstrates broad conservation of disease-associated T cell populations across human and murine synovial tissue, apart from Tregs, which exhibit high PD-1 expression only in the murine model, and identifies PD-1 as a therapeutically relevant target for the development of pathogenic T-cell directed CAR-T cell therapy in RA.

### PD-1 targeting CAR-T cells undergo antigen-dependent activation and selectively deplete PD-1⁺ T cells

To selectively target disease-associated synovial T cells expressing high levels of PD-1, we incorporated the sequence of a previously characterized high-affinity monoclonal antibody against the extracellular domain of murine PD-1^51,52^. This sequence was reformatted into a single-chain variable fragment (scFv) and integrated into a third-generation chimeric antigen receptor (CAR) construct. The CAR was delivered using a retroviral backbone, and the intracellular signaling domain included tandem co-stimulatory modules; CD28 and 4-1BB followed by the CD3ζ signaling chain (Fig. 2a).

**Figure 2.**
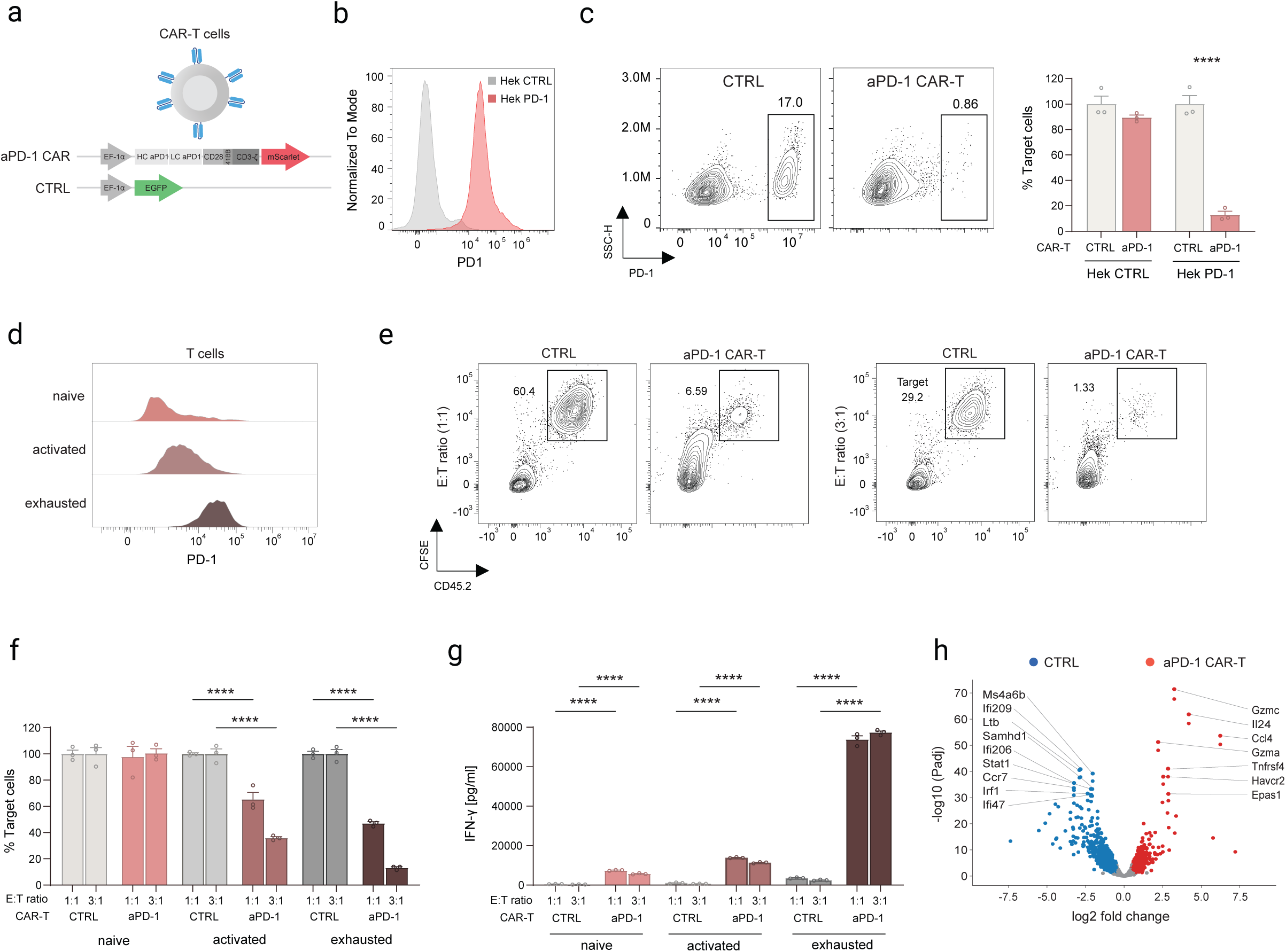
aPD-1 CAR-T cells mediate potent antigen-specific activation and selective clearance of PD-1^+^ cells. (a) Schematic illustration of aPD-1 CAR-T and control GFP vector (CTRL). (b) representative flow cytometry analysis for PD-1 expression on HEK293T cells over-expressed (OE) murine *Pdcd1* (Hek PD-1) versus HEK293T cells only (Hek CTRL). (c) Left. Representative flow cytometry analysis of Hek CTRL and Hek PD-1 percentages following 24h co-cultured with aPD-1 CAR-T and T cells transduced with control GFP vector (CTRL transduced T cells) at E:T ratio of 5:1. Right. Bar plots represent relative proportion of Hek PD-1 or Hek CTRL cells after 24h co-cultured with aPD-1 CAR-T cells and T cells transduced with CTRL transduced T cells at E:T ratio of 5:1. Data are presented as mean ± SD (n = 3). Statistical significance was assessed by Two-way ANOVA (****p < 0.0001). (d) Representative flow cytometry analysis of PD-1 expression on naïve, activated, or exhausted bulk T cells. (e) Representative flow cytometry analysis of bulk T cell percentages following 24h co-cultured with mouse aPD-1 CAR-T or CTRL transduced T cells at E:T ratio of 1:1 and 3:1. (f) Bar plots represent relative proportion of naïve, activated, or exhausted bulk T cells after 24h co-cultured with mouse aPD-1 CAR-T and CTRL transduced T cells at E: T ratio of 1:1 and 3:1. Data are presented as mean ± SD (n = 3). (****p < 0.0001). (g) Mouse IFN-γ ELISA assay following 24h co-cultured of aPD-1 CAR-T and CTRL transduced T cells with naïve, activated, or exhausted bulk T cells at E: T ratio of 1:1 and 3:1. Data are presented as mean ± SD (n = 3). Statistical significance was assessed by Two-way ANOVA (**** p < 0.0001). (h) Volcano plot of DEGs in aPD-1 CAR-T cells compared to CTRL transduced T cells in vitro. Genes with adjusted P < 0.05 and |log₂FC| > 1 were highlighted.

Bulk T cells were transduced with the CAR construct with high efficiency (70-90%, Supplementary Fig. 4a). To assess the target specificity of PD-1 targeting CAR-T cells, we co-cultured them with either parental HEK293T cells (Hek CTRL) or HEK293T cells overexpressing PD-1 (Hek PD-1, Fig. 2b). Cytotoxicity of aPD-1 CAR-T cells against PD-1 expressing target cells was assessed by flow cytometry (Fig. 2c). aPD-1 CAR-T cells efficiently and selectively depleted Hek PD-1 cells, resulting in ∼80% reduction in target cell survival compared to parental Hek CTRL cells or Hek PD-1 cells co-cultured with control T cells transduced with an EF-1α-GFP control vector (CTRL, Fig. 2a and Fig. 2c). Furthermore, activation-induced IFN-γ secretion into the co-culture supernatant was quantified by ELISA (Supplementary Fig. 4b). Activated aPD-1 CAR-T cells showed increased IFN-γ secretion when co-cultured with Hek PD-1 cells, but not with parental cells or when Hek PD-1 cells co-cultured with control-transduced T cells, confirming PD-1-specific, antigen-dependent CAR-T cell activation (Supplementary Fig. 4b).

To evaluate the ability of aPD-1 CAR-T cells to eliminate their physiological targets, we assessed their cytotoxic activity against murine bulk T cells isolated from the spleen. aPD-1 CAR-T cells were co-cultured with either naïve bulk T cells, once-activated bulk T cells, or bulk T cells activated through three successive rounds (hereafter referred to as exhausted), which expressed low, intermediate, and high levels of PD-1, respectively (Fig. 2d). After 24 hours, aPD-1 CAR-T cells at 3:1 and 1:1 effector-to-target (E:T) ratios efficiently and selectively depleted PD-1^high^ exhausted T cells, resulting in an approximately 90% reduction in target cell survival at the highest E:T ratio. In contrast, an approximately 60% reduction was observed in PD-1^intermediate^ cells, while PD-1^low^ cells and PD-1 expressing T cells co-cultured with control T cells remained unaffected (Fig. 2e-f). In addition to target cell killing, aPD-1 CAR-T cells exhibited robust, dose-dependent IFN-γ secretion in response to PD-1 expressing targets, correlating with both target-cell frequency and PD-1 expression level of target T cells (Fig. 2g).

Consistent with these findings, aPD-1 CAR-T cells showed markedly increased cytotoxicity, reaching ∼70% at a 3:1 ratio when co-cultured with T cells transduced with a PD-1 expression vector (Supplementary Fig. 4c-d). In contrast, no cytotoxicity was observed when control-vector transduced T cells were incubated with aPD-1 CAR-T cells, confirming antigen-specific activity (Supplementary Fig. 4c-d). No reduction in viability or evidence of fratricide among aPD-1 CAR-T cells was observed during these killing assays (Supplementary Fig. 4e). To validate CAR-T cell activation and cytotoxicity at the molecular level following co-incubation with PD-1^high^ target T cells, we sorted CAR-T cells after co-culture with or without their targets at varying effector-to-target (E:T) ratios for scRNA-seq (Fig. 2h). Compared with CAR-T cells alone or control-transduced T cells, antigen-exposed CAR-T cells upregulated genes associated with activation, effector differentiation, cytokine production, and cytotoxicity (e.g., *Gzmc*, *Gzma*, *Tnfrsf4*, *Ccl4*). In contrast, control-transduced T cells remained in a less differentiated/memory-like state, showing minimal non-specific activation (Fig. 2h). Our results demonstrate that aPD-1 CAR-T cells are specifically activated by PD-1⁺ T cells *in vitro* and mount a potent, selective cytotoxic response, effectively eliminating PD-1 expressing target cells without loss of viability or evidence of fratricide.

### aPD-1 CAR-T cells selectively target synovial disease-associated T cells and suppress arthritis

To enable mechanistic dissection of *in vivo* T cell responses in arthritis following aPD-1 CAR-T therapy, we first employed the mBSA-induced arthritis model. Arthritis was induced in the knee and ankle joints of the left hind limb, while the corresponding joints on the right side were left untreated as controls. Three days after arthritis induction, at the peak of joint inflammation, mice were intravenously infused with either 10 million CD45.1⁺ congenic aPD-1 CAR-T cells, or with T cells transduced with EF-1α-GFP control vector (CTRL). During the re-challenge phase (characterized by greater expansion of Tph cells relative to the initial phase, Fig. 1j), aPD-1 CAR-T cell therapy significantly reduced arthritis severity in both knee and ankle joints by day 9 after induction compared to controls (Fig. 3a-b, Supplementary Fig. 5a). Similar efficacy was observed when aPD-1 CAR-T cell administered at disease induction, attenuating both the initial response and subsequent re-challenge (Supplementary Fig. 5b). aPD-1 CAR-T therapy was well tolerated, with no apparent adverse effects or measurable weight loss observed in treated mice.

**Figure 3.**
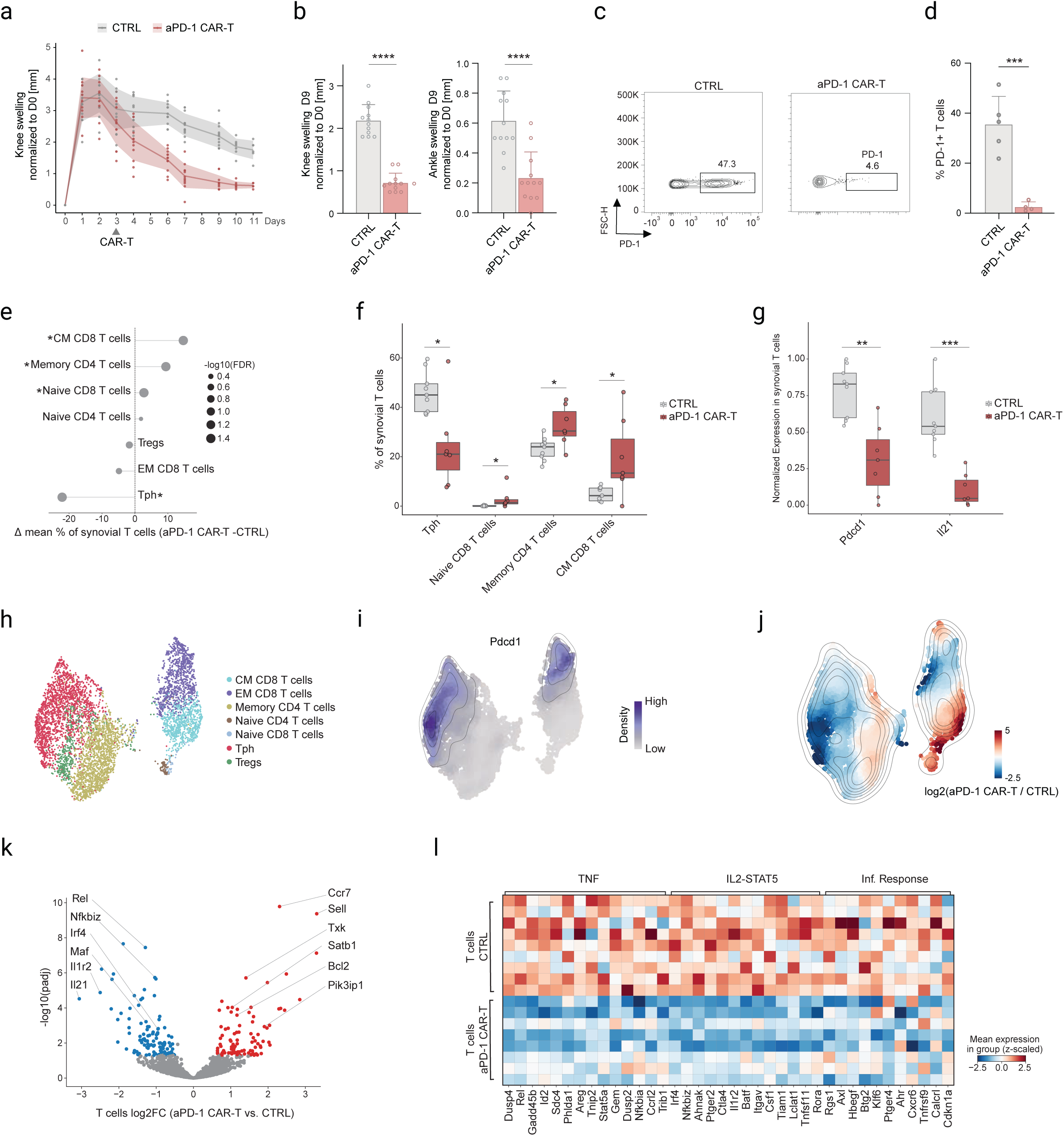
aPD-1 CAR-T cells selectively deplete PD-1^+^ disease-associated T cells in vivo, and significantly mitigate arthritis in two murine models. (a) *In vivo* efficacy of PD-1 CAR-T cell therapy in mice following arthritis induction during re-challenge phase using the mBSA arthritis model. Arthritis severity was monitored by caliper measurement of knee joint swelling (y-axis, mm) over time following arthritis induction (x-axis, days). CAR-T cells were administered on day 3, after disease establishment. Data are presented for two groups: control-transduced T cells (CTRL), and aPD-1 CAR-T cells. n=12-13 mice/group. (b) Summary of joint swelling during re-challenge phase. Bar graphs depict mean knee (left) and ankle (right) joints swelling (mm above baseline diameter) in the two experimental groups (control, aPD-1 CAR-T cells) at day 9 following arthritis induction. One-way ANOVA (**** p < 0.0001). (c) Representative flow cytometry analysis of PD-1^+^T cells in synovial tissue of mBSA-induced arthritic joints following aPD-1 CAR-T therapy (right) or CTRL transduced T cells administration (left) at day 9 post-CAR or CTRL injection. (d) Bar graphs depicting the frequencies of PD-1^+^ T cell in arthritis joints following aPD-1 CAR-T or CTRL transduced T cells therapies, shown per individual experiment. One-way ANOVA (***p < 0.001). (e) Difference in the average percentage of T cells in the synovium of aPD-1 CAR-T treated mice (n=7) compared to controls (n=9). Dot size represents the statistical significance of the difference. (* P < 0.05). (f) Boxplot showing the percentage of significantly different T subsets from total synovial T cells of arthritis mice under aPD-1 CAR-T treatment (n=7) compared to CTRL transduced T cells group (n=9). Each point represents a mouse. (* P < 0.05). (g) Boxplot showing normalized expression of Pdcd1 and Il21 in synovial T cells from CD45^+^ gate of aPD-1 CAR-T treated mice (n=7) compared to CTRL transduced T cells group (n=9). (** P < 0.01), (*** P < 0.001). (h) Integrated T cells UMAP, with graph-based clustering of 5,828 T cells from the synovium of 22 arthritis mice, down sampled for an equal number of cells per gate in each condition. (i) Density plot showing distribution of Pdcd1 normalized expression on the synovial T cells embedding, down sampled for an equal number of cells per gate in each condition. (j) Density plot showing the delta log_2_FC of synovial T cells embedding density of T cells under aPD-1 CAR-T treatment and CTRL, down sampled for an equal number of cells per gate in each condition. (k) Volcano plot of DEGs in synovial T cells under aPD-1 CAR-T compared to CTRL transduced T cells therapy. Genes with adjusted P < 0.05 and |log₂FC| > 0.5 were highlighted. (l) Heatmap showing expression of DEGs from specific pathways in synovial T cells pseudo bulked per mouse across conditions.

To investigate the *in vivo* mode of action of the aPD-1 CAR-T construct and its impact on PD-1⁺ disease-associated T cells, mice were injected with either aPD-1 CAR-T or control transduced T cells 3 days after arthritis induction (during the re-challenge phase). Eight days after CAR-T cell injection, synovial tissues (knee and ankle) and spleens were harvested, dissociated, and analyzed by flow cytometry and scRNA-seq. Congenic aPD-1 CAR-T cells accumulated in arthritic joints and the spleen but were markedly reduced in contralateral non-arthritic joints (Supplementary Fig. 5c), while control-transduced T cells were restricted to the spleen. scRNA-seq analysis of *in vivo*-derived synovial CAR-T cells post-therapy demonstrated a cytotoxic, proliferative, and activated phenotype (e.g., *Gzmb*, *Prf1*, *Mki67*, *Top2a*), whereas naïve synovial T cells expressed genes associated with a less activated state (e.g., *Sell*, *Il7r*, *Tcf7*; Supplementary Fig. 5d).

Flow cytometric analysis of synovial tissue demonstrated significant depletion of synovial PD-1⁺ T cells in mice treated with aPD-1 CAR-T cells relative to the control group (Fig. 3c-d). scRNA-seq analysis of synovial T cells following aPD-1 CAR-T cell therapy showed an overall reduction in synovial T cells following treatment (Supplementary Fig. 5e-f), with preferential depletion of host PD-1⁺ T cells, most prominently within the PD-1^high^ disease-associated Tph subset and a trend toward decreased PD-1^high^ EM CD8^+^ T cells (Fig. 3e-f, Fig. 3h-j, Supplementary Fig. 5f). A modest effect was also observed in synovial Tregs, which express

lower PD-1 levels (Fig. 3h-j). We further observed a significant decrease in *Pdcd1* and *Il21* expression in synovial T cells following aPD-1 CAR-T therapy, consistent with reduced Tph cell frequency, a population that is a key source of IL-21 supporting local B cell responses (Fig. 3g). In the spleen of aPD-1 CAR-T treated mice, no significant reduction in total T cell numbers was observed (Supplementary Fig. 5g). However, Tregs were selectively reduced, with a trend toward decreases of PD-1^high^ EM CD8^+^ T cells compared to control group (Supplementary Fig. 5h-k).

Interestingly, following aPD-1 CAR-T cell therapy, both spleen and synovium were repopulated with naïve T cells, suggesting broad remodeling of the T cell compartment across tissues, while the synovium remained enriched for memory T cell subsets (Fig. 3e-f. Fig. 3h-j, Supplementary Fig. 5h-k). Residual synovial T cells after aPD-1 CAR-T cell therapy showed reduced expression of *Il21*, *Il1r2*, *Maf*, *Irf4*, *and Nfkbiz,* alongside increased expression of *Ccr7*, *Sell*, and *Bcl2*, consistent with a shift toward a less activated, central memory-like phenotype (Fig. 3k). In parallel, pseudo-bulk analysis revealed pronounced attenuation of TNF, IL-2-STAT5, and inflammatory response pathways following CAR-T cell therapy (Fig. 3l, Supplementary Fig. 5l). Together, these data indicate that CAR-T cell therapy not only depletes pathogenic populations but also promotes reprogramming of the synovial T cell landscape toward a less inflammatory and more regulated state.

To directly compare aPD-1 CAR T-cell therapy with antibody-based approaches, we generated depleting (IgG2) and Fc-silent blocking (IgG1-D265A) aPD-1 antibodies sharing the same antigen-binding (Fab) domain as the CAR-T construct. Mice were treated with 250 μg of each antibody intravenously, aPD-1 CAR or control-transduced T cells on day 3 after arthritis induction (Supplementary Fig. 5m). Neither antibody reduced disease severity, whereas aPD-1 CAR-T therapy significantly attenuated arthritis (Supplementary Fig. 5m). By day 11 following treatment, aPD-1 CAR-T cells induced robust depletion of PD-1^+^ T cell populations. In comparison, the aPD-1 depleting antibody produced modest reductions in PD-1^+^ T cells in both joint and spleen, while the aPD-1 blocking antibody showed no reduction and, in some cases, was associated with increased PD-1^+^ T cell frequencies (Supplementary Fig. 5-n). These findings highlight the enhanced efficacy of CAR-T therapy at the tested dose, particularly at the tissue level. To extend and corroborate these findings while incorporating the B cell component of disease, we employed the spontaneous systemic polyarthritis K/BxN model, which recapitulates key features of human RA, including autoreactive T cell-dependent B cell activation and autoantibody production^53^. This enabled validation of T cell-associated programs identified in the mBSA model while also allowing analysis of pathogenic T-B cell interactions. K/BxN arthritis is driven by a defined adaptive immune response in which autoreactive CD4⁺ T cells support B cells to induce pathogenic autoantibody production, triggering downstream innate effector mechanisms in the joint. In this setting, PD-1^high^ T follicular helper (Tfh) cells in lymphoid tissues and peripheral helper (Tph) cells in inflamed joints are key drivers of autoreactive B cell responses^54^.

Synovial tissue of K/BxN mice before the onset of clinical disease (pre-disease phase: 3-3.5 weeks) and after clinical disease development (4.5-5.5 weeks) showed an increase in PD-1⁺ T cells in the joints following disease development, with no significant changes in B cell percentages (Supplementary Fig. 6a-e). Adoptive transfer of 1.8 million CAR-T cells generated from splenic T cells of pre-disease K/BxN mice into stage-matched syngeneic recipients markedly prevented or attenuated disease development compared with control-transduced T cells from age-matched mice or untreated controls (Supplementary Fig. 6f). This was accompanied by a significant reduction in both PD-1⁺ T-cell abundance and PD-1 expression, as well as decreased myeloid cell infiltration in the joints, but not in the spleen, relative to mice treated with control-transduced T cells, or untreated mice (Supplementary Fig. 6g-j).

Reduced PD-1⁺ T cell abundance in K/BxN synovium was accompanied by decreased *Ccr1* and *Litaf* expression in synovial-resident T cells, consistent with reduced inflammatory cell recruitment and attenuation of TNF-driven inflammation (Supplementary Fig. 6m). In parallel, splenic T cells following CAR-T cell therapy showed reduced expression of *Icos, Irf4*, and *Batf* (Supplementary Fig. 6n). Given the central role of ICOS signaling in sustaining Tfh/Tph differentiation and B cell helper function via PI3K-AKT-dependent induction of *Irf4* and *Batf* ^55^, this reduction is consistent with attenuated T-B cell interactions. Concomitantly, splenic T cells demonstrated increased *Eomes* expression, alongside upregulation of *Sh2b3* and *Dusp6*, consistent with enhanced negative regulation of TCR and cytokine signaling pathways (Supplementary Fig. 6n). Overall, aPD-1 CAR Therapy dampens ICOS-associated T-cell gene expression signatures and T-B cell interactions in the spleen, while enhancing pathways involved in the negative regulation of TCR and cytokine signaling. Similarly, following CAR-T cell therapy, splenic B cells exhibited coordinated downregulation of activation and inflammatory signaling pathways (*Stat4, Il6st, Cd69, and Tlr7*), alongside upregulation of regulatory and homeostatic pathways (*Grn, Sgk1, Abcg1*, *Pou2f2,* Supplementary Fig. 6o). Consistent with these cell-intrinsic changes, ligand-receptor analysis of splenic B-T interactions showed reduced *Icos* signaling following aPD-1 CAR-T therapy versus controls, alongside increased *Tgfb, Lair2*, and *Klrd1* signaling (Supplementary Fig. 6p). This shift suggests rewiring from co-stimulatory, disease-promoting interactions toward a more regulated intercellular signaling landscape.

Together, these data reveal a coordinated remodeling of adaptive immunity following aPD-1 CAR-T therapy, in which both T and B cell compartments are reprogrammed toward a restrained, non-pathogenic state, with diminished B cell helper function, while preserving core immune functionality. Across both mBSA and K/BxN arthritis model, aPD-1 CAR-T therapy reduced PD-1^+^ synovial T cells, attenuated inflammatory signaling and reshaped T-B cell communication, supporting a broad immunomodulatory mechanism of disease control.

### Engineering context-dependent regulatory elements for synovium selective CAR-T function

aPD-1 CAR-T cells effectively ameliorated joint inflammation and were well tolerated, with no overt macroscopic toxicity. However, constitutive CAR expression can promote tonic, antigen-independent signaling, limiting spatial specificity, particularly in autoimmune settings where the target antigen is also expressed on non-pathogenic cells in peripheral tissues. This raises the risk of off-target cytotoxicity and disruption of immune homeostasis. To address this limitation, we engineered a regulated CAR system that couple’s receptor expression to endogenous inflammatory transcriptional programs identified from transcriptomic data, enabling context-dependent activation. This design aims to restrict CAR activity to inflamed synovial tissue while limiting activation in peripheral compartments, thereby reducing collateral effects.

To achieve tissue-selective CAR activity, we performed scRNA-seq of aPD-1 CAR-T cells from both inflamed synovium and spleen to define tissue-specific activation state and identify transcription factors circuits enriched in diseased tissue, which were subsequently used to engineer synthetic regulatory elements driving context-dependent CAR expression (Fig. 4a-b). Consistent with enhanced local activation, expression of *Nr4a1/2* was increased *in vivo* in both aPD-1 CAR-T and endogenous T cells within the inflamed joint compared with splenic isolated aPD-1 CAR-T cells (Fig. 4b-c). Because Nr4a family members are induced downstream of sustained TCR/CAR signaling, particularly in antigen-engaged T cells within chronically inflamed joints, we leveraged their binding motifs to regulate a distal CAR enhancer (Fig. 4d). The NR4A2 motifs were assembled into synthetic regulatory elements comprising four or six tandem repeats organized as transcription factor binding clusters, thereby generating DNA biosensors that drive gene expression in response to CAR-T cell activation (Fig. 4d). As candidate regulatory elements controlling CAR expression, we employed both homotypic NR4A2 motifs and heterotypic bipartite NFAT-NR4A2 motifs designed to recapitulate cooperative transcription factors binding within the TCR/CAR activation program. NFAT, a canonical proximal-response regulatory motif, marks acute antigen engagement and is therefore associated with broader off-target activation, whereas NR4A2 reflects more distal, sustained activation, thereby restricting CAR activity to chronically stimulated, antigen-engaged T cells in inflamed tissue^56,57^.

**Figure 4.**
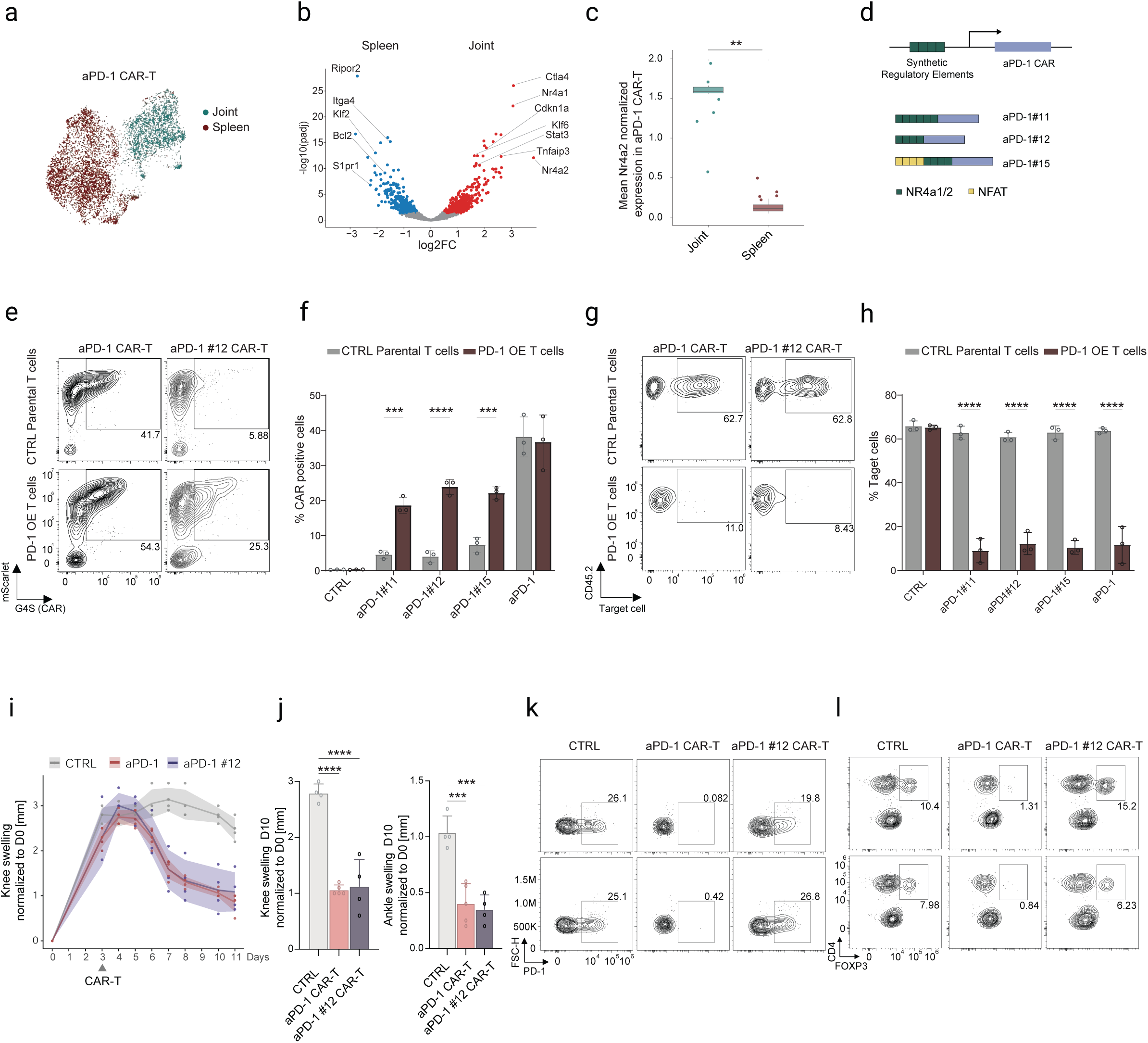
Synovium-specific regulatory elements enable controlled aPD-1 CAR activation with reduced off-target effects. (a) Integrated UMAP of aPD-1 CAR-T cells isolated from the spleen and synovium of six arthritis mice with tissue annotations. (b) Volcano plot of DEGs in aPD-1 CAR-T from synovium compared to the spleen of arthritis mice. Genes with adjusted P < 0.05 and |log₂FC| > 0.5 were highlighted. (c) Boxplot showing normalized expression of Nr4a2 in aPD-1 CAR-T of synovium and spleen of arthritis mice. (** P < 0.01). (d) Schematic illustration of aPD-1 CAR-T design incorporating synthetic joint tissue-specific regulatory elements proximal to the CAR promotor. (e) Representative flow cytometry analysis of G4S and mScarlet expression in constitutive CD45.1^+^ aPD-1 CAR-T cells versus CD45.1^+^ regulated aPD-1#12 CAR-T cells following 24 h of incubation with CD45.2^+^ PD-1 overexpressing (OE) bulk T cells or CD45.2^+^ control parental T cells transduced with an empty vector at E:T ratio of 3:1. (f) Bar plots summarizing G4S staining by flow cytometry in constitutive aPD-1 CAR-T cells, various regulated aPD-1 CAR-T cells, and control-transduced T cells (CTRL) 24 h after incubation with target cells, including PD-1 overexpressing (OE) bulk T cells or control parental T cells transduced with an empty vector, at an E:T ratio of 3:1. Data are presented as mean ± SEM. (***P <0.001), (****P <0.0001). (g) Representative flow cytometry analysis of residual CD45.2^+^ PD-1 overexpressing (O.E.) bulk T cells or CD45.2^+^ control parental T cells transduced with an empty vector, following 24 h of incubation with constitutive CD45.1^+^ aPD-1 CAR-T cells or CD45.1^+^ regulated aPD-1 #12 CAR-T cells at an E:T ratio of 3:1. (h) Bar plots summarizing staining by flow cytometry for bulk T CD45.2^+^ target cells overexpressing (O.E.) PD-1 or CD45.2^+^ parental T cells transduced with an empty vector, following 24 h of incubation with constitutive CD45.1^+^ aPD-1 CAR-T cells, CD45.1^+^ regulated aPD-1 #12 CAR-T cells, or CD45.1^+^ controlled-transduced T cells (CTRL) at an E:T ratio of 3:1. Data are presented as mean ± SEM, (****P<0.0001). (i) *In vivo* efficacy of PD-1 CAR-T, regulated aPD-1 #12 CAR, or controlled-transduced T cells (CTRL) therapies in mice following arthritis induction during re-challenge phase using the mBSA arthritis model. Arthritis severity was monitored by caliper measurement of knee joint swelling (y-axis, mm) over time following arthritis induction (x-axis, days). CAR-T cells or CTRL were administered on day 3, after disease establishment. Data is presented for three groups: control-transduced T cells (CTRL), aPD-1 CAR-T cells, or aPD-1#12 CAR-T cells. n=5 mice/group. (j) Summary of joint swelling during re-challenge phase. Bar graphs depict mean knee and ankle joint swelling (mm above baseline diameter) in the three experimental groups (CTRL, aPD-1 CAR-T cells, aPD-1#12 CAR-T cells) at day 10 following arthritis induction. One-way ANOVA (***p <0.001), (**** p < 0.0001). (k-l) Representative flow cytometry analysis of PD-1⁺ T cells (k) and Tregs (l) from the spleen of mBSA-induced arthritic mice at day 10 after injection of aPD-1 CAR-T cells, aPD-1 #12 CAR-T cells, or control-transduced T cells (CTRL).

To assess biosensor specificity and functional activity, CAR-T cells were transduced with plasmids encoding biosensor variants in which regulatory elements were positioned upstream of the aPD-1 CAR promoter, along with a constitutive mScarlet reporter to monitor transduction efficiency. Surface CAR expression was quantified by flow cytometry using an anti-G4S antibody recognizing the extracellular Gly₄Ser (G4S) linker, enabling specific and antigen-independent detection of CAR expressed at the cell surface. aPD-1 CAR-T cells transduced with different regulatory elements were co-cultured with either PD-1 overexpressing (OE) target T cells or control (CTRL) parental T cells transduced with an empty vector. Biosensor activation and CAR functionality were assessed by flow cytometry via CAR (G4S) staining and by measuring target cell killing, respectively (Fig. 4e-h, Supplementary Fig. 7a-b). Flow cytometry analysis demonstrated constitutive G4S staining in cells expressing the constitutive aPD-1 CAR, irrespective of ligand expression on target cells (Fig. 4e-f, Supplementary Fig. 7a). In contrast, G4S expression in the regulated CARs was detected only following co-culture with PD-1 OE target T cells, but not with control cells, and occurred in a response element-dependent manner (Fig. 4e-f, Supplementary Fig. 7a). No G4S expression was observed in control transduced T cells, confirming a direct link between CAR-mediated signaling and biosensor-driven gene expression (Supplementary Fig. 7a). Importantly, although all regulated designs reduced basal CAR expression relative to the constitutive CAR, they differed substantially in signal to background behavior. aPD-1#11 and aPD-1#12 showed the lowest basal G4S expression on control parental T cells, whereas aPD-1#15, which incorporates the NFAT-containing bipartite element, exhibited significantly higher background activity. Upon co-culture with PD-1 OE target T cells, aPD-1#12 showed the strongest inducible CAR expression among the regulated constructs, thereby providing the most favorable balance between low basal activity and robust target-induced activation (Supplementary Fig. 7a).

Similar killing of target cells overexpressing PD-1 was observed across the different regulated CAR constructs and the constitutive CAR, whereas no killing was observed with control-transduced T cells (Fig. 4g-h, Supplementary Fig. 7b). We further validated PD-1 target-specific cytotoxicity by co-incubating aPD-1 CAR-T cells harboring different regulatory elements with Hek cells expressing luciferase (mock) or overexpressing the target PD-1 antigen and luciferase, followed by a luciferase-based killing assay (Supplementary Fig. 7c). All aPD-1 regulated CAR constructs demonstrated specific and effective killing of PD-1 expressing target cells compared to control-transduced T cells and at levels comparable to constitutively expressed CAR (Supplementary Fig. 7c). Thus, while all regulated constructs preserved target-dependent cytotoxic function, the principal differences between them lay in basal and inducible CAR expression rather than in endpoint killing. To assess the *in vivo* activation of regulated CARs, including their therapeutic efficacy and ability to limit peripheral off-target effects, we evaluated aPD-1 CAR-T cell function in a murine mBSA arthritis model. CAR-T cells expressing three regulatory constructs (aPD-1#11, aPD-1#12, aPD-1#15), a constitutive aPD-1 CAR, or control-transduced T cells were administered three days after arthritis re-challenge (Fig. 4i-j4J, Supplementary Fig. 7d). Consistent with its favorable *in vitro* signal to background profile, the regulatory construct aPD-1#12 CAR improved disease severity, with efficacy comparable to the constitutive aPD-1 CAR, whereas the other regulated constructs showed more modest benefit (Supplementary Fig. 7d). Consistent with context-restricted activity, aPD-1#12 CAR resulted in reduced depletion of splenic PD-1 expressing cells, specifically FOXP3^+^ Tregs compared to the constitutive CAR, indicating diminished off-target cytotoxicity (Fig. 4k-l). Collectively, these findings show that aPD-1#12 CAR maintains therapeutic efficacy while enhancing selectivity and identifying it as the optimal regulatory construct based on its low basal activity, strong inducibility, preserved cytotoxicity, and improved in vivo selectivity specifically to splenic Tregs.

### sTNFRii-armored aPD-1 CAR-T cells further enhance tissue-restorative immune remodeling

After establishing a CAR construct that effectively depletes PD-1 expressing cells and markedly attenuates arthritis, we aimed to further enhance its therapeutic impact by engineering an additional cassette encoding an anti-inflammatory cargo. This strategy was designed to counteract the pro-inflammatory influence of bystander cells within the inflammatory niche, potentially further amplifying its therapeutic benefit. We engineered a murine sTNFRii cargo (Fig. 5a), representing the soluble form of the TNF receptor that binds and neutralizes TNF-α, thereby competitively inhibiting its interaction with membrane-bound receptors. This blockade prevents activation of downstream pro-inflammatory signaling cascades, effectively attenuating inflammation and limiting tissue damage^58–61^. In clinical applications, recombinant sTNFRii, most notably in the form of the fusion protein etanercept, has been employed as a key therapy for RA^61^. To validate sTNFRii expression and secretion, murine T cells were transduced with sTNFRii, aPD-1 CAR-T, a GFP control vector, or co-transduced with both the aPD-1 CAR and sTNFRii (aPD-1 sTNFRii CAR-T, Fig. 5a). High levels of sTNFRii were detected in the supernatants of T cells transduced with sTNFRii alone or engineered as sTNFRii-secreting CAR-T cells, but not in those transduced with the aPD-1 CAR or a control vector (Fig. 5a). The bioactivity of sTNFRii secreted by aPD-1 CAR-T cells was validated using HEK293T cells expressing TNF receptors and an NF-κB inducible SEAP reporter to detect TNF-α activity. Inhibition of the reporter signal by aPD-1 sTNFRii CAR-T cells, but not by aPD-1 CAR- T cells alone or control-transduced T cells, confirmed that the engineered cargo effectively neutralizes TNF-α (Fig. 5b).

**Figure 5.**
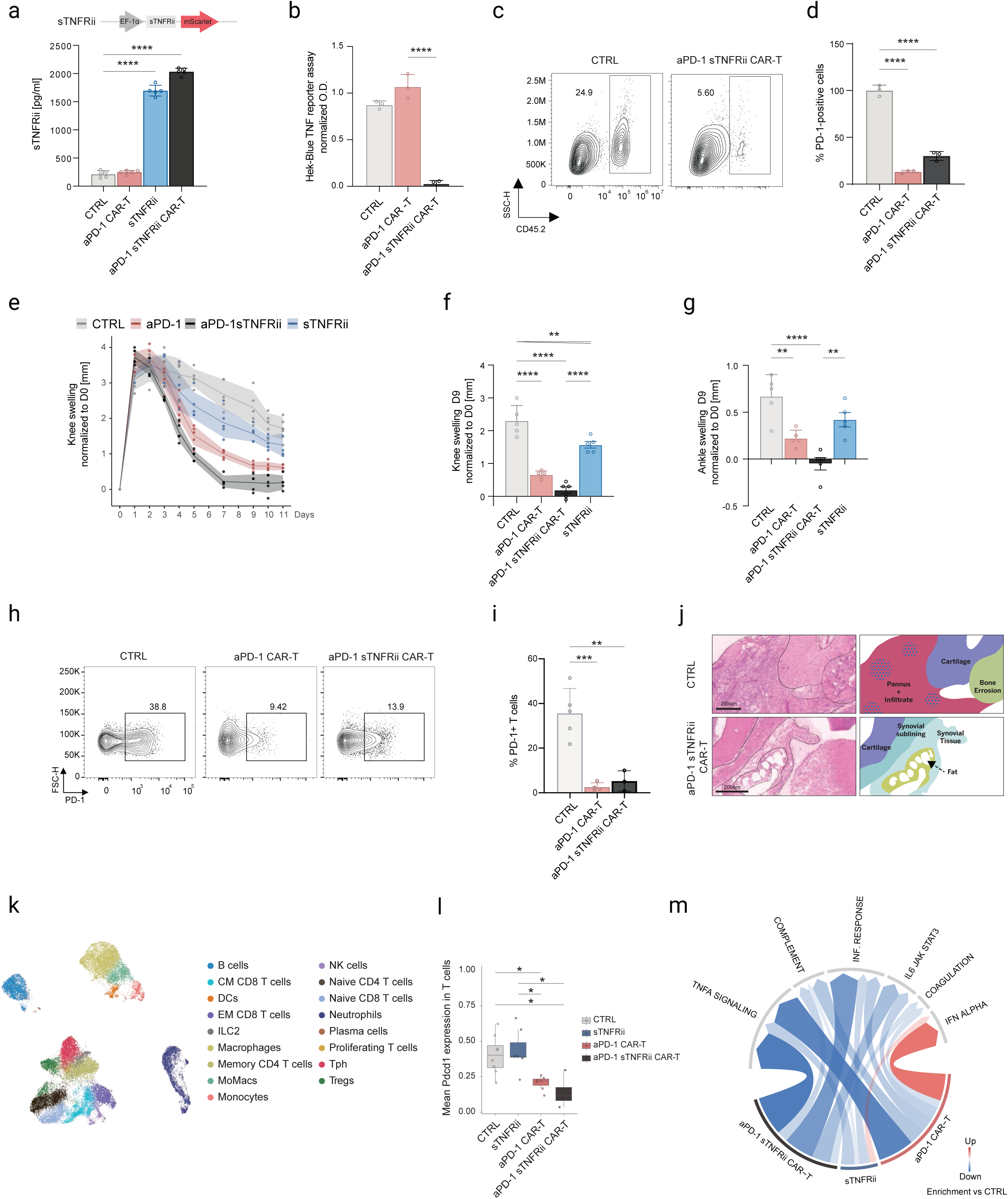
sTNFRii-secreting aPD-1 CAR-T cells therapy mitigates arthritis and reshapes the synovial immune microenvironment. (a) Quantification of soluble TNF receptor ii (sTNFRii) secretion was performed by ELISA on supernatants collected from T cells transduced with sTNFRii cargo alone, aPD-1 CAR, or both aPD-1 CAR and sTNFRii. Bound TNFRii was detected using a biotinylated anti-TNFRii antibody followed by streptavidin-HRP, and absorbance was measured at 450 nm. Data are presented as mean ± SEM (pg/ml) from triplicate wells and are representative of n = 5 independent experiments One-way ANOVA (*** p < 0.0001). (b) HEK-Blue reporter assay for detection of secreted activity upon cargo stimulation. HEK-Blue™ reporter cells (5 × 10⁴ cells/well) were incubated with the indicated cargo and TNF-α (0.1 ng/ml) for 12 h. Cell supernatants were collected and analyzed using QUANTI-Blue™ detection reagent, and absorbance was measured at 620-655 nm using a plate reader. Data are presented as mean ± SEM of triplicate wells and are representative of n = 2 independent experiments. Statistical significance was determined by one-way ANOVA (****P < 0.0001). (c) Representative flow cytometry analysis of Hek CTRL and Hek PD-1 percentages following 24h co-cultured with aPD-1 sTNFRii CAR-T cells or with control GFP vector (CTRL) at E: T ratio of 5:1. (d) Barplots represent relative proportion of Hek CTRL or Hek PD-1 after 24h co-cultured with aPD-1 CAR-T, aPD-1 sTNFRii CAR-T cells, and control transduced-T cells (CTRL) at E: T ratio of 5:1. Data are presented as mean ± SD (n = 3). (**** P < 0.0001). (e) *In vivo* efficacy of bulk T cells transduced with sTNFRii cargo only, versus PD-1 CAR-T or PD-1 sTNFRii CAR-T cell therapies versus controlled-transduced T cells (CTRL), in mice following arthritis induction during re-challenge phase using the mBSA arthritis model. Arthritis severity was monitored by caliper measurement of knee joint swelling (y-axis, mm) over time following arthritis induction (x-axis, days). Cargo only, CTRL and CAR-T cells therapies were administered on day 3, after disease establishment. Data are presented for four groups: sTNFRii, aPD-1 CAR-T cells, aPD-1 sTNFRii CAR-T cells, CTRL. n=5 mice/group. (f-g) Summary of knee (f), or ankle (g) joint swelling during re-challenge phase. Bar graphs depict mean joint swelling (mm above baseline diameter) in the four experimental groups (sTNFRii, aPD-1 CAR-T cells, aPD-1 sTNFRii CAR-T cells, CTRL) at day 9 following arthritis induction. One-way ANOVA (**** p < 0.0001), (h) Representative FACS analysis of PD-1^+^ T cell percentages in synovial tissue of mBSA-induced arthritic joints at day 9 post aPD-1 CAR-T cells, aPD-1 sTNFRii CAR-T cells or controlled-transduced T cells (CTRL) injection. (i) Bar graphs depicting the frequencies of PD-1^+^ T cell in arthritis joints following PD-1, aPD-1-sTNFRii CAR-T or control-transduced T cell therapies (CTRL), shown per individual experiment. One-way ANOVA (**p < 0.01). (***p < 0.001). (j) Histopathology of arthritic joints following CAR-T cell therapy. Representative H&E-stained sections of decalcified, cryo-fixed joints (n=3) from C57BL/6 mice with arthritis, collected 8 days after treatment with aPD-1 CAR-T cells or control-transduced T cells (CTRL) during the re-challenge phase. Scale bar, 200 µm. (k) Integrated immune cells UMAP, with graph-based clustering of 44,869 immune cells from the synovium and spleen of arthritis mice across treatments. (l) Boxplot showing normalized expression of Pdcd1 in synovial T cells across treatments - CTRL (n=8), sTNFRii (n=7), aPD-1 CAR-T (n=7), aPD-1 sTNFRii CAR-T (n=4). *P<0.05. (m) Chord diagram showing the normalized enrichment scores of the Hallmark defined pathways among myeloid cells across different treatments. Red and blue lines indicate upregulation and downregulation of the modules, respectively. Color coding corresponds to the treatment.

To determine whether addition of the cargo altered CAR function, we performed a killing assay and co-cultured aPD-1 sTNFRii CAR-T cells, and aPD-1 CAR-T cells with either parental Hek control cells or Hek PD-1^+^ cells (Fig. 5c). Both aPD-1 sTNFRii CAR-T cells and aPD-1 CAR-T cells efficiently and selectively depleted Hek PD-1^+^ cells (60-75% reduction in survival), and produced high levels of IFN-γ when co-cultured with Hek PD-1^+^cells, but not with parental Hek control cells or when Hek PD-1^+^cells were co-cultured with control-transduced T cells (Fig. 5c-d, Supplementary Fig. 8a). Taken together, these findings establish aPD-1 sTNFRii CAR-T cells as a dual-function therapeutic, capable of selectively depleting PD-1 expressing target cells, while concurrently attenuating TNFα-driven inflammation in the local environment. To assess the *in vivo* effect of sTNFRii secretion by aPD-1 CAR-T cells, four groups of mice were treated 3 days after arthritis mBSA re-challenge with aPD-1 CAR-T cells, sTNFRii-transduced T cells, aPD-1 sTNFRii CAR-T cells, or control-transduced T cells (Fig. 5e). Eleven days later, joints were harvested and dissociated for analysis. While both aPD-1 CAR-T therapies markedly improved the arthritic phenotype relative to the control-transduced and cargo-only groups, near-complete clinical resolution was observed only in mice treated with aPD-1 sTNFRii CAR-T cells (Fig. 5e-g).

Flow cytometry analysis confirmed the presence of aPD-1 CAR-T cells, with or without cargo, in both the inflamed joint and spleen, whereas T cells transduced with cargo alone or control transduced T cells were detected only in the spleen. We further observed depletion of PD-1⁺ T cells and a reduction in PD-1-expressing in T cells in the synovium following treatment with aPD-1 CAR-T cells, irrespective of cargo, but not with control-transduced T cells or T cells transduced with cargo alone as confirmed by both flow cytometry and scRNA-seq (Fig. 5h-i, Fig. 5k-l, Supplementary Fig. 8b). These changes align with histopathological evidence of disease resolution, including reduced inflammation and bone erosion (Fig. 5j). To elucidate the impact of secreted sTNFRii cargo on the synovial microenvironment, we performed comprehensive scRNA-seq analyses of synovial immune cells before and after treatment with the different constructs, with a focus on the myeloid compartment, a key driver of TNFα, IL-1, and IL-6-mediated inflammation in RA, where synovial macrophages promote leukocyte recruitment, stromal activation, matrix degradation, and osteoclastogenesis, leading to cartilage destruction and bone erosion^20^.

T cells transduced with sTNFRii alone induced a limited, systemic anti-inflammatory effect compared with controls, reflected by modest reductions in *Il6*, *Cxcl1/2*, and *Inhba*, without targeted modulation of the synovial microenvironment (Supplementary Fig. 8c-e). In contrast, aPD-1 CAR- T therapy alone more effectively suppressed inflammation, reducing TNFα, IL-1, and IL-6-driven pathways, angiogenesis signaling, and chemokine-mediated leukocyte recruitment, but remained associated with an interferon-skewed, activated phenotype compared with aPD-1 sTNFRii CAR-T cells (Fig. 5m, Supplementary Fig. 8c-d), which may contribute to cytokine release syndrome in patients (CRS)^62^. Notably, sTNFRii-secreting CAR-T cells exerted the strongest overall effect, achieving the most pronounced suppression of inflammatory programs, including interferon signaling, and angiogenic programs, surpassing both aPD-1 CAR-T and cargo-only treatments. This was accompanied by upregulation of *Tgfb1, Lyve1, Cx3cr1, Siglec1,* and *Ccl24* in synovial myeloid cells, consistent with a shift toward a regulatory, tissue-repairing, and pro-resolving phenotype (Supplementary Fig. 8c-d).

In line with this, both aPD-1 CAR-T and aPD-1 sTNFRii CAR-T cells localized within the joint exhibited significantly higher expression of *CD69* and the interferon-stimulated gene *RTP4* compared to circulating cargo-only transduced T cells, consistent with tissue retention and local activation within an interferon-rich inflammatory microenvironment (Supplementary Fig. 8e). Together, these data define a clear hierarchy of therapeutic efficacy: circulating T cells transduced with sTNFRii alone remain undetectable in the synovium and yield only modest systemic effects; aPD-1 CAR-T cells that infiltrate and activate within the joint suppress local inflammatory circuits and leukocyte recruitment, albeit with increased interferon signaling; and aPD-1 sTNFRii CAR-T cells most effectively inhibit pathogenic pathways while promoting a tissue-restorative state.

## Discussion

Despite major advances in DMARDs and biologics development, a substantial subset of RA patients remains difficult to treat or poly-refractory^63–65^, highlighting the need for therapies that more effectively eliminate disease-sustaining immune populations. The persistence of long-lived disease-associated T and B cell clones within the synovium, an immune-privileged niche that allows their survival and re-expansion and thereby promotes relapse, represents a major barrier to sustained disease control. Our findings suggest that selective targeting of disease-associated synovial T cells may provide such a strategy by disrupting core pathogenic circuits within the inflamed tissue niche.

CAR-T cell therapy, which has been transformative in hematologic malignancies, is increasingly being explored as a potential modality for autoimmune disease. Early clinical reports of CD19- and BCMA-targeted CAR-T cells suggest the possibility of durable remissions in B cell-mediated autoimmune disorders^11,12,17,18^. However, these approaches may be associated with broad depletion of B cell populations, incomplete targeting of pathogenic clones, and an increased risk of infection, underscoring the need for next-generation designs with enhanced specificity and safety.

Using multi-omics profiling of RA patient synovium and a murine arthritis model, we functionally define Tph cells as key drivers of disease pathology and identify PD-1 expression as a marker of these active clonotypes. aPD-1 CAR-T cells eliminated PD-1⁺ T cells both *in vitro* and *in vivo*, reduced the frequency of pathogenic PD-1⁺ Tph cells, and ameliorated arthritis severity, alongside increased representation of naïve and central memory T cells. These findings parallel observations of immune reconstitution following anti-CD19 or BCMA CAR-T therapies in B cell-mediated autoimmune diseases and hematological malignancies and reflect a shift toward a less differentiated and more homeostatically regulated T cell compartment^11,66^.

As a distinct subset in RA, Tph cells exhibit stem-like properties, including self-renewal and persistent replenishment, enabling the maintenance of a reservoir of pathogenic T cells within inflamed tissue^26,40^. Concurrently, they provide critical help to B cells through IL-21 and CXCL13, driving activation, differentiation, and autoantibody production. The ability of our CAR-T approach to deplete these cells points to a therapeutic strategy that targets a central pathogenic mechanism that sustains both T cell- and B cell-driven disease processes. By targeting these interconnected arms of the adaptive immune response, this approach may enable deeper and more durable disease control than strategies focused on a single compartment. PD1^+^ Tph cells are also implicated in RA-associated interstitial lung disease^67^, and in other autoimmune disorders, such as systemic lupus erythematosus and primary Sjögren’s syndrome^68,69^, suggesting that targeting this axis may extend therapeutic benefit across different autoimmune diseases.

Single-cell profiling of aPD-1 CAR-T cells from the inflamed synovium and spleen of arthritic mice identified Nr4a1/2 as synovium-enriched transcription factors that could be leveraged to confer disease-restricted control of CAR activity. We therefore designed and evaluated an Nr4a2-based regulatory circuit that couples CAR expression to the inflamed tissue state and limits splenic depletion of PD-1-expressing cells, including tissue-resident Tregs. Collectively, these experiments provide a conceptual basis for integrating tissue-responsive regulatory circuits into CAR-T design, enabling more precise disease-targeted activity and potentially safer cellular therapies for autoimmune disease.

Building on the therapeutic efficacy of aPD-1 CAR-T cell therapy and its ability to home to the target organ without lymphodepletion or IL-2 preconditioning, we engineered sTNFRii-secreting aPD-1-targeting CAR-T cells that combine selective depletion of disease-associated T cells with local TNF-α neutralization. This combinatorial approach not only attenuated inflammation but also reshaped the synovial microenvironment by restoring anti-inflammatory transcriptional programs and suppressing pro-inflammatory myeloid signatures. These changes were associated with preservation of synovial tissue integrity in murine arthritis models and may further support enhanced CAR-T cell activity and persistence. The strategy of combining selective depletion of disease-associated T cells with local TNF-α neutralization may offer a more refined and potentially safer therapeutic approach, with the potential to attenuate cytokine release syndrome (CRS) and immune effector cell-associated neurotoxicity syndrome (ICANS).

Our findings define a cooperative therapeutic paradigm that integrates site-specific and selective depletion of disease-associated T cells with localized immunomodulation. By simultaneously targeting pathogenic T cell populations and reshaping the inflamed tissue microenvironment, this approach addresses key limitations of current RA therapies, which often incompletely eliminate disease-driving cells or fail to durably reprogram tissue inflammation. Targeting disease-associated T cells within the affected tissue may complement existing CAR-T and bispecific antibody strategies in autoimmune diseases. While B cell-directed CAR-T therapies have shown substantial efficacy, tissue-resident pathogenic T cell populations may also contribute to disease persistence. Selective targeting of these cells *in situ* may help modulate local inflammatory circuits and T-B cell interactions, potentially improving precision and enabling more durable disease control. Finally, our work establishes the conceptual basis for multi-omics-guided CAR-T engineering across T cell or mixed T/B cell-mediated autoimmune diseases, with the potential to enable durable immune reset and sustained remission. Beyond RA, this disease-localized CAR-T platform provides a roadmap for extending this strategy to other autoimmune diseases driven by pathogenic T cells or coordinated T/B cell programs.

## Acknowledgments

We would like to thank Itai Raveh for the artwork. I.A. is an Eden and Steven Romick Professorial Chair, supported by the HHMI International Scholar Award, funded by the European Union ERC advanced grant (no. 101055341-TROJAN-Cell), and by the Deutsche Forschungsgemeinschaft (DFG, German Research Foundation) -Project-ID 259373024 - TRR 167, and the Israel Science Foundation grant no. 1944/22, the Helen and Martin Kimmel awards for innovative investigation, Israel Science Foundation Precision Medicine Program (IPMP) 607/20, Dwek Institute for Cancer Therapy Research, Moross Integrated Cancer Center, Morris Kahn Institute for Human Immunology, Swiss Society Institute for Cancer Prevention Research, Elsie and Marvin Dekelboum Family Foundation, Lotte and John Hecht Memorial Foundation and the Schwartz Reisman Collaborative Science Program and the Weizmann center for Immunotherapy. T.S.P is supported by postdoctoral fellowship of the Azrieli Foundation and T.S.P and K.F. by the faculty postdoctoral excellence fellowship of the Weizmann Institute of Science.

The results published here are in part based on data obtained from the ARK Portal (https://arkportal.synapse.org/). This work was supported by the Accelerating Medicines Partnership® Rheumatoid Arthritis and Systemic Lupus Erythematosus (AMP® RA/SLE) Program. AMP® is a public-private partnership (AbbVie Inc., Arthritis Foundation, Bristol-Myers Squibb Company, Foundation for the National Institutes of Health, GlaxoSmithKline, Janssen Research and Development, LLC, Lupus Foundation of America, Lupus Research Alliance, Merck & Co., Inc., National Institute of Allergy and Infectious Diseases, National Institute of Arthritis and Musculoskeletal and Skin Diseases, Pfizer Inc., Rheumatology Research Foundation, Sanofi and Takeda Pharmaceuticals International, Inc.) created to develop new ways of identifying and validating promising biological targets for diagnostics and drug development Funding was provided through grants from the National Institutes of Health (UH2-AR067676, UH2-AR067677, UH2-AR067679, UH2-AR067681, UH2-AR067685, UH2- AR067688, UH2-AR067689, UH2-AR067690, UH2-AR067691, UH2-AR067694, and UM2- AR067678).

## Resource availability

Lead contact

Further information and requests for resources and reagents should be directed to and will be fulfilled by the lead contact, Ido Amit.

## Materials availability

Correspondence and requests for materials should be addressed to Ido Amit. All biological materials can be obtained from the corresponding authors following reasonable requests.

## Data and code availability

Single-cell RNA-seq data will be deposited in the Gene Expression Omnibus and will be made publicly available as of the date of publication. Any additional information required to re-analyze the data reported in this paper is available from the lead contact upon request. Osteoarthritis control dataset was taken from the Gene Expression Omnibus deposit with the accession number GSE248453.

This Study uses human TCR-seq data from Argyriou A* Marc H. Wadsworth II*; Nat Commun. 2022 Jul 13;13(1):4046. doi: 10.1038/s41467-022-31519-6. Human PBMC data in this study are taken from CellxGene Discover resource https://cellxgene.cziscience.com/collections/e1a9ca56-f2ee-435d-980a-4f49ab7a952b. 16 The results published here are in whole or in part based on data obtained from the ARK Portal (https://arkportal.synapse.org/). This work was supported by the Accelerating Medicines Partnership® Rheumatoid Arthritis and Systemic Lupus Erythematosus (AMP® RA/SLE) Program. AMP® is a public-private partnership (AbbVie Inc., Arthritis Foundation, Bristol-Myers Squibb Company, Foundation for the National Institutes of Health, GlaxoSmithKline, Janssen Research and Development, LLC, Lupus Foundation of America, Lupus Research Alliance, Merck & Co., Inc., National Institute of Allergy and Infectious Diseases, National Institute of Arthritis and Musculoskeletal and Skin Diseases, Pfizer Inc., Rheumatology Research Foundation, Sanofi and Takeda Pharmaceuticals International, Inc.) created to develop new ways of identifying and validating promising biological targets for diagnostics and drug development Funding was provided through grants from the National Institutes of Health (UH2-AR067676, UH2-AR067677, UH2-AR067679, UH2-AR067681, UH2-AR067685, UH2- AR067688, UH2-AR067689, UH2-AR067690, UH2-AR067691, UH2-AR067694, and UM2- AR067678).

## Author contributions

C.G. calibrated and developed experimental protocols; conceptualized, designed, performed, and analyzed experiments, interpreted results, wrote and reviewed the manuscript ; L.R. performed all bioinformatic analyses; analyzed experiments and interpreted results and wrote and reviewed the manuscript; R.S.E. calibrated and developed experimental protocols; conceptualized, designed, performed, and analyzed experiments and interpreted results; R.S contributed to calibration of experimental protocols and performed in vitro experiments; R.A. contributed to the development of cargo secretion regulatory mechanisms. E.R designed and generated overexpressing PD-1 HEK293T cells and T cells. M.N.V.L. advised with reporter assay experiments. K.X. and E.D. advised the computational pipeline. R.S., M.B.Y. and G.Y. contributed to the CAR design and performed experiments. M.Z. advised on in vitro experiments. E.D. and K.M. conducted the scRNA-seq library preparation, sequencing, and read alignment. Y.K. performed in vivo experiments. K.A. performed histological experiments. Y.N., H.P., M.L., A.B.G. reviewed the manuscript. S.K.E. and R.T helped perform in vitro experiments. T.S.P. developed experimental protocols; conceptualized, designed, performed, and analyzed experimental results, supervised analyses, wrote and reviewed the manuscript K.F. developed experimental protocols; conceptualized, designed, performed, and analyzed experiments and interpreted results, supervised the study, wrote and reviewed the manuscript. I.A. developed experimental protocols, mentored and directed the project, conceptualized and designed experiments, interpreted results, and wrote the manuscript.

## Declaration of generative AI and AI-assisted technologies in the writing process

During the preparation of this work, the authors used ChatGPT in order to improve writing clarity. After using this tool, the authors reviewed and edited the content as needed and take full responsibility for the content of the publication.

## Author declaration

The authors have submitted a patent application pertaining to the findings described in this manuscript.

## Materials and Methods

### Mouse

Experimental cohorts included male and female wild type C57BL/6 (CD45.2) or K/BxN mice (3-8 weeks old). C57BL/6 (CD45.1, strain #002014), and KRN mice were obtained from The Jackson Laboratory. Using C57BL/6 CD45.1 mice allows tracking of engineered T cells derived from CD45.1 donors after transfer into CD45.2 recipients. KRN TCR transgenic mice were crossed with NOD mice to generate K/BxN mice according to established protocols^53^. All animals were housed under specific pathogen-free conditions in the Weizmann Institute animal facility, maintained on a 12-hour light/12-hour dark cycle with libitum access to food and water. All procedures were performed in accordance with protocols approved by the Institutional Animal Care and Use Committee.

### Mouse Arthritis Models

mBSA-induced arthritis was established according to the protocol^70^: To induce arthritis, 8–10-week-old C57BL/6 mice were immunized subcutaneously with 100 μg of methylated bovine serum albumin (mBSA; 2 mg/mL in sterile water) emulsified in complete Freund’s adjuvant (CFA; 1:1, vol/vol), administered in 100 μL to the right hind flank. Simultaneously, 160 ng of pertussis toxin (1.6 μg/mL in PBS) was injected intraperitoneally in 100 μL. A booster injection of 100 μg mBSA/CFA was administered subcutaneously to the left hind flank 7 days later. On day 0 (21 days after the initial immunization), arthritis was induced by intra-articular injection of 30-50 μL mBSA (10 mg/mL in sterile water) into the left ankle and knee (into the infrapatellar space) joints, respectively, under anesthesia with ketamine-xylazine. The right knee and ankle received sterile water as a control. Joint swelling was monitored using a digital micrometer.

K/BxN mice were generated by crossing KRN TCR transgenic mice with NOD mice^53^. Progeny expressing both the KRN transgene and I-A^g7 spontaneously develops arthritis driven by anti-glucose-6-phosphate isomerase autoantibodies. Arthritis usually became clearly established at ∼4-5 weeks of age and was assessed by clinical scoring (0-4 per paw) and ankle thickness measurements.

### Cell lines

HEK293T cells engineered to overexpress mouse *Pdcd1* and/or luciferase were maintained in DMEM (Thermo Fisher, 11965084) supplemented with 10% fetal calf serum (FCS), 100 U/mL penicillin, and 100 µg/mL streptomycin. Plat-E cells were used for retroviral production and cultured in DMEM supplemented with 10% FBS and 1% penicillin-streptomycin. For selection, Plat-E cells were maintained in 1 µg/mL puromycin and 10 µg/mL blasticidin.

### Cloning of CAR Constructs Plasmid design and synthesis

To generate the aPD-1 CAR, the heavy and light chains of the antigen recognition domains of aPD-1 antibody were utilized as binding domains. The aPD-1 CAR construct was constructed by assembling a single-chain variable fragment (scFv) in a VH-linker-VL orientation, followed by the CD28 and 4-1BB transmembrane and cytoplasmic domains, and the CD3ζ cytoplasmic domain. The complete CAR design was cloned into the self-inactivating (SIN) retroviral backbone (Addgene, plasmid Cat. No. 153417). The CAR sequence was fused to a T2A-mScarlet reporter for visualization and tracking. The sTNFRii cargo and the GFP were cloned into the same retroviral backbone (self-inactivating (SIN) retroviral backbone (Addgene, Cat. No. 153417)). The complete plasmid sequences were ordered as double-stranded genes from GenScript. For *Pdcd1* overexpression, HEK293T cells were transfected with a pLEX-based plasmid encoding murine Pdcd1 (Addgene plasmid Cat. No. 162033). To express CAR-PD1 under environmental sensors, we designed a transcriptional element composed of DNA motifs that fit the interface of Nr4a1 (UNIPROT:P12813) and Nr4a2 (UNIPROT:Q06219) DNA binding domains. The DNA binding zinc finger domains of both Nr4a1 and Nr4a2 share 95% homology and both bind to the classical hexamer sequence AGGTCA, that is recognized by all transcription factors of the nuclear receptor family. To optimize the sequence for Nr4a1 and Nr4a2 binding, we extended the 5’ site of the hexamer with an AT-rich motif^71^ with two extra adenines (AAAGTCA) to favor Nr4a1 or Nr4a2 binding as a homodimer on the positive strand, which is joined to a palindromic sequence of the hexamer and further extended with A and T bases (TGACCTAT). To avoid steric hinderance and allow for equal motif usage on both strands, both motifs were separated by an inert 5bp sequence that does not bind to any residues on the DNA binding domain of both TFs. To modulate activity of the synthetic element (AACTGAAAGGTCAAACTGTGACCTAT), the Nr4a binding motifs were joined together as 4 repeats (CAR-PD1 #11) or 6 repeats (CAR-PD1 #12), with each repeat being separated by the inert sequence for the same purpose. The sequences generated were then predicted for motif binding using MEME Suite FIMO^72^. The combination of the NR4a1/2 repeated motifs with the of the NFAT TF (Car-PD1 #15) was generated as previously described^73^.

### Retrovirus production

For virus production, Plat-E cells were seeded at 70-80% confluency in DMEM media without antibiotics and transfected using Lipofectamine 2000 (11668027, Thermo Fisher) with the retroviral vector of interest along with the packaging plasmid pCL-Eco (Addgene plasmid Cat. No. 12371). Six to eight hours after transfection, the medium was replaced with fresh DMEM containing 10% FBS. Viral supernatants were collected at 48 hours post-transfection, filtered through a 0.45 µm filter, and used immediately or snap frozen and stored at −80°C.

### Generation of aPD-1 CAR-T cells

For transduction, 24-well non-tissue culture-treated plates were coated with RetroNectin (25 µg/ml in PBS) at 0.5 ml per well for 2 hours at room temperature or overnight at 4 °C. Plates were blocked with PBS containing 2% BSA for 30 minutes at room temperature and then washed with HBSS supplemented with 2.5% 1 M HEPES. Viral supernatants were added directly onto the coated plates for single or double transductions and centrifuged at 2,000 g for 2 hours at 32 °C to allow virus adsorption. In parallel, primary murine T cells were isolated from spleens using a Pan T cell isolation kit (Miltenyi Biotec 130-095-130) activated with anti-CD3/CD28- antibodies (Ultra-LEAF™ Purified anti-mouse CD3 [BLG-100340], Ultra-

LEAF™ Purified anti-mouse CD28 [BLG-102116]), and cultured overnight in lymphocyte medium supplemented with 10 ng/ml recombinant murine IL-2 (PeproTech 212-12-100). Following RetroNectin loading, viral supernatants were removed, and 1×10⁶ activated T cells in 1 ml of lymphocyte medium containing IL-2 were added per well by gently pipetting along the wall to avoid disturbing the RetroNectin coating. Plates were centrifuged at 350 g for 20 minutes at 32 °C and incubated overnight at 37 °C. The following day, cells were collected and transferred to new tissue culture-treated plates. IL-7 (10 ng/ml, Rhenium 217-17-100) and IL-15 (10 ng/ml), PreptoTech 210-15-100) were added to the culture medium starting on day 2 post-transduction to support further expansion.

### Generation of aPD-1 blocking or depleting antibodies

To generate depleting versus blocking antibodies, we used an IgG2a isotype as the depleting format in the wild-type mouse system. In parallel, an Fc-silent version (mIgG1-D265A) was included as a control to distinguish between cell depletion and PD-1 blocking activity. The antibodies are based on the anti-PD-1 clone RMP1-14. The following constructs were generated: RMP1-14 IgG1-D265A heavy chain (460 amino acids) paired with the RMP1-14 light chain (238 amino acids), and RMP1-14 IgG2a heavy chain (471 amino acids) paired with the RMP1-14 light chain (238 amino acids). The depleting antibody design was based on the approach described in^51^. All antibody genes were synthesized and generated by GenScrip.

### HEK-Blue TNF-α Reporter Assay

HEK-Blue™ TNF-α cells were engineered from the human embryonic kidney HEK293 cell line to detect bioactive murine tumor necrosis factor-alpha (TNF-α) by monitoring the activation of the NF-κB and AP-1 pathways. HEK-Blue™ TNF-α cells were generated by stable transfection with the genes encoding for the murine TNF-α receptor (TNFR1 and TNFR2 chains), as well as an NF-κB/AP-1-inducible SEAP secreted embryonic alkaline phosphatase (SEAP) reporter. The binding of TNF-α to its receptor triggers a signaling cascade leading to NF-κB/AP1 activation and the subsequent production of SEAP. This can be readily assessed in the supernatant using QUANTI-Blue™ Solution, a SEAP detection reagent. For functional assessment, HEK Blue™ TNF-α cells were incubated overnight at 37°C with serial tenfold dilutions of supernatant collected from CAR-T cells 24 hours after antigen exposure and 0.1 ng/ml of mouse TNF-α. SEAP levels in the culture supernatants were measured using QUANTI-Blue™ Solution and quantified at 630-650 nm with a SpectraMax Plus spectrophotometer (Molecular Devices).

### Measurement of mouse IFN-γ and Cargo secretion

IFN-γ levels were quantified using ELISA MAX™ Deluxe Sets for mouse IFN-γ (BLG567 430804) from BioLegend. 96-well ELISA plates were coated overnight at 4°C with 0.2 µg/well of capture antibody diluted in PBS. Plates were then washed three times with 0.05% Tween-20 in PBS, blocked with 1% BSA in PBS for 1 hour at room temperature (RT), and washed again. Serum samples were added to the coated wells and incubated for 2 hours at RT. After washing, detection antibodies were applied, followed by incubation with streptavidin-HRP for 20 minutes at RT. Plates were again washed and developed with substrate solution F for 20 minutes in the dark at RT, then stopped by adding 2N sulfuric acid (DY994, R&D). Optical density was measured at 450 nm with a 570 nm reference using a SpectraMax Plus spectrophotometer (Molecular Devices).

Soluble TNFRii (Tnfrsf1b) secretion from CAR-T cells transduced with the sTNFRii plasmid was quantified using the Boster Picokine® Mouse Tnfrsf1b Pre-Coated ELISA (Enzyme-Linked Immunosorbent Assay) kit-a solid-phase immunoassay specially designed to measure Mouse Tnfrsf1b with a 96-well strip plate that is pre-coated with antibody specific for Tnfrsf1b. The detection antibody is a biotinylated antibody specific to Tnfrsf1b, and the kit includes Mouse Tnfrsf1b protein as standards. To measure Mouse Tnfrsf1b, we added standards and samples to the wells, added the biotinylated detection antibody, washed the wells with TBS buffer, and added Avidin-Biotin-Peroxidase Complex (ABC-HRP). After washing away, the unbounded ABC-HRP with TBS buffer we added TMB. The absorbance of the yellow product at 450nm is linearly proportional to Mouse Tnfrsf1b in the sample.

### Synovial tissue processing

Synovial cells were isolated from murine joints by careful dissection of periarticular tissues. Following euthanasia, hind limbs were disinfected, and skin, tendons, ligaments and muscles were removed to expose the knee or ankle joint. The synovial membrane and adjacent soft tissues (e.g., infrapatellar fat pad) were dissected without breaching the bone to avoid bone marrow contamination. Tissues were enzymatically digested in DMEM without serum containing 0.3mg/mL Liberase, 2U/ml Dispase, and 1 mg/mL DNase I at 37°C for 75 minutes with gentle agitation. Digestion was quenched with cold medium containing 10% FBS, and the suspension was filtered through a 70 µm strainer. Cells were pelleted by centrifugation and subjected to red blood cell lysis using ACK lysis buffer for 5 minutes on ice. Lysis was stopped with cold medium, and cells were washed, pelleted again, and resuspended in appropriate buffer for downstream applications.

### Isolation of splenic T cells

Spleens were harvested under sterile conditions and mechanically dissociated by gentle disruption through a 70-µm cell strainer (BD Biosciences) to obtain single-cell suspensions. Red blood cells were removed by treatment with RBC lysis buffer (eBioscience, Thermo Fisher) according to the manufacturer’s instructions. Following lysis, cells were washed twice with cold phosphate-buffered saline (PBS) and centrifuged at 300×g for 10 minutes. The resulting splenocyte suspensions were used directly for downstream applications, or T cells were further isolated by negative selection using the Pan T Cell Isolation Kit II, mouse (Miltenyi Biotec, Bergisch Gladbach, Germany) following the manufacturer’s protocol.

### Exhaustion protocol

Bulk T cells were subjected to a repeated stimulation protocol consisting of three sequential anti-CD3 stimulations every 48 hours, with anti-CD28 co-stimulation provided only during the first stimulation. Control cells received CD3 and CD28 co-stimulation at the first stimulation only. Following the final activation, cells were analyzed by flow cytometry for *Pdcd1* expression and viability.

### Flow Cytometry Staining and Analysis

Single-cell suspensions were prepared as previously described. Cells isolated from synovial tissues were stained with anti-CD45.1 PE/Cy7 (clone A20, 110730, BD), anti-CD45.1 APC/Cy7 (clone A20, 560579, BD), anti-CD45.2 PE (clone 104, 109808, BD), anti-CD45.2 APC (clone 104, 109814, BD), anti-CD90.2 FITC (clone 53-2.1, 140304, BD), anti-CD69 PE/Cy7 (clone H1.2F3, 104511, BD), anti-TCR b chain APC/Cy7 (clone H57-597, 109219, BD), anti-CD279 APC (clone 29F.1A12, 135209, BD), anti-CD11b (clone M1/70, 101257, BioLegend), anti-CD279 APC/Cy7 (clone 29F.1A12, 135223, BD).Anti-G4S PE Linker (clone E7O2V, 38907S, Cell Signaling), anti-G4S AF700 linker (cloneE7O2V, 40107S, Cell Signaling), anti-CD4 (clone RM4-5, 100548, BioLegend), anti-CD3 (clone 17A2, 100222, BioLegend), anti-CD25 (clone PC61, 102007, BioLegend), anti-FOXP3 (clone R16-715, 567373, BD).We used Anti-CD16/CD32 TruStain FcX™ (clone 93; BioLegend, #101320, #630) to block FcγRIII/II receptors and prevent non-specific binding, incubating for 10 minutes on ice. Live dead staining: Zombie aqua BLG-423102 or DAPI (4’,6-diamidino-2- phenylindole). Cells were analyzed on a BD FACS Symphony 6 flow cytometer and FlowJo software (version 10). For killing assay of bulk T cells by CAR-T, target cells were stained with CellTrace ^TM^ CFSE/Far red cell proliferation kits (C34554, C34572 Invitrogen) according to the protocol.

### Single cell library preparation

scRNA-seq libraries were prepared using SPID-seq^50^, a modified version of the MARS-seq protocol^49^. Briefly, polyadenylated mRNA from single cells sorted into 384-well plates was captured and barcoded during reverse transcription into cDNA. cDNA from each plate was then pooled, fragmented, and amplified to generate sequencing-ready libraries for Illumina NovaSeq X Plus. Quality control was conducted for each plate-derived library.

Read Alignment Pooled libraries were sequenced on an Illumina NovaSeq X Plus platform at a depth of 10,000-50,000 reads per cell. Reads with identical unique molecular identifiers (UMIs) collapsed to represent individual RNA molecules. Batch quality was validated by confirming low cross-cell contamination (<3%), based on the frequency of spurious UMIs in empty wells. Alignment was performed using the MARS-seq2.0 pipeline: low-quality reads were removed, and remaining reads were aligned to the mouse genome (mm10) using HISAT (v0.1.6), excluding multimapping reads. Only exonic reads were counted, based on UCSC gene annotations. UMI uniqueness was confirmed within a 3 kb window. For overlapping exons from distinct genes on the same strand, reads were assigned to a merged gene ID combining both gene symbols.

### Preprocessing and analysis of SPID-seq data

Quality control and preprocessing were performed using Scanpy^74^ ( v1.10.4) for all mice datasets. Cells with fewer than 300 counts, more than 15,000 counts, fewer than 300 detected genes, or over 20%mitochondrial reads were excluded. Ribosomal and mitochondrial genes were filtered out prior to dimensionality reduction. The top 5,000 most variable genes were selected. Principal component analysis (PCA) was performed using 50 components. A neighborhood graph was then computed using 50 principal components and 10 nearest neighbors and projected into a 2D space using UMAP for visualization (min_dist=0.3), and clustering was performed using the Leiden algorithm. Cell populations were manually annotated based on the expression of canonical marker genes. Differential gene expression analysis was performed using pyDESEQ2^75^ and Decoupler^76^ using Wald test. Genes were filtered by adjusted p < 0.05 and absolute log fold-change > 0.5 unless mentioned otherwise.

For cell-type proportions calculations, mice with more than 2 cell-type proportions that were not in the lower 25% or more than 75% across all mice were excluded. Proportions and expression differences were calculated using the Mann-Whitney U test, and p-values were corrected using Benjamini-Hochberg FDR correction.

### Human synovial and PBMC single-cell atlas construction and analysis

Atlas samples were collected from different available datasets^21,24,37,39,44^. Quality control and preprocessing were performed using the Python package Scanpy^74^. Cells with fewer than 500 counts, more than 50,000 counts, or more than 20% of reads mapped to mitochondrial genes were excluded. Cells classified as doublets by Scrublet^77^ were removed. The data was integrated using scVI^45^. The scVI model was trained for 400 epochs, using 1,000 highly variable genes for the T cells and 5,000 highly variable genes for the immune compartment. Batch correction was performed using the sample name as batch key, and single cell sequencing technology and the study name as categorical covariates. Cell type annotations were performed by unsupervised graph-based clustering using the Leiden algorithm, followed by manual annotation based on canonical markers.

### Mouse single-cell TCR seq analysis

Single-cell TCR repertoire sequencing was performed on isolated T cells from synovial tissue of 11 mice (22 joint)8 days following arthritis induction using the mBSA-induced arthritis model. Raw data were processed with Scanpy^74^ and Scirpy^78^ to identify and quantify clonotypes. Downstream analyses, including clonotype frequency, clonal expansion and overlap between samples, were performed using Scirpy^78^. Integration of the TCR sequences and SPID sequences was performed using scVI^45^. 5,000 highly variable genes were used for the integration, and the single cell technology was used as ‘batch key’. The model was trained for 400 epochs. For the two samples from the TCR sequencing, we down sampled for an equal number of cells.

### Human synovial and PBMC single-cell TCR and BCR seq ligand-receptor interaction analysis

Samples were downloaded from^37^. Quality control and preprocessing of scRNA data were performed using Scanpy^74^. Cells with fewer than 500 counts, more than 25,000 counts, more than 20% of reads mapped to mitochondrial genes or more than 50% of reads mapped to ribosomal genes were excluded. The RNA data was integrated using scVI^45^, with 5,000 highly\ variable genes and patient as batch key. The model was trained for 400 epochs. Cell type annotations were performed by unsupervised graph-based clustering using the Leiden algorithm, followed by manual annotation based on canonical markers. TCR and BCR data were processed using Scirpy^78^ and integrated with the RNA data using Mudata^79^. T and B cells with multichain, orphan VDJ, orphan VJ, or ambiguous chain pairings were removed. Circular bar plots for T and B cells were generated by counting unique clones shared in more than 1 cell per cell type in each tissue. A circos plot of ligand-target network between synovial expanded T cells as senders and synovial expanded B cells as receivers was generated by dividing the respective T cells into PD1^+^ and PD1^-^ using raw UMI counts. Genes expressed in at least 10% of cells in each group were taken for senders, and differential genes with |log2FC| > 0.25 in the expanded synovial B cells were taken for receivers. Nichenet was employed to analyze and predict cell-cell interactions from sc RNA seq^80^.

### K/BxN single-cell based splenic T-B interactions

MultiNichenet was employed to analyze and predict cell-cell interactions from K/BxN mice sc RNA seq^80^. Scanpy object was converted to SingleCellExperiment object^74^ for downstream analysis. Interactions analyses were conducted for aPD-1 CAR-T treated mice and for untreated mice separately. For each analysis, T cells and B cells from the spleen were considered as both senders and receivers. Genes expressed in at least 5% of the cells, at least 50% of all samples, at least 10 cells, |log2FC| > 0.5, and P < 0.05 were considered in each group.

### Treatment Density Calculation

Populations abundance for aPD1 CAR-T treated T cells was calculated by down sampling of treated and untreated T cells from TCRβ sorted plates and was projected using the Scanpy function embedding density^74^.

### Myeloid cells pathways enrichment analysis and modules generation

Monocytes, macrophages and MoMacs were extracted from CD45^+^ sorted plates, across synovium samples of untreated, aPD-1 CAR-T or aPD-1 sTNFRii CAR-T treated mice. The preprocessing step was done as mentioned above. Differential expression analysis was conducted on pseudo bulk myeloid cells per mouse using Decoupler^76^ and pyDESEQ2^81^. Pathway enrichment analysis was done for each treatment compared to control, using the Hallmark curated pathways database. P values were then corrected using Benjamini-Hochberg FDR correction, and non-significant pathways were excluded, in addition to non-inflammatory related pathways. For inflammation-associated modules, specific DEGs were considered, and a heatmap was generated. Modules expression differences among treatments were calculated using the Mann-Whitney U test, and p-values were corrected using Benjamini-Hochberg FDR correction.

### Statistical analysis

For cell-types proportions, expression and scores comparisons across conditions, two-sided Mann-Whitney U test was conducted. If more than one hypothesis was tested, FDR Benjamini-Hochberg correction was applied for the p-values. Statistical significance was denoted as follows: p < 0.05: *, p < 0.01: **, p < 0.001: ***, p<0.0001: ****

## Supplementary Figures

**Figure S1.**
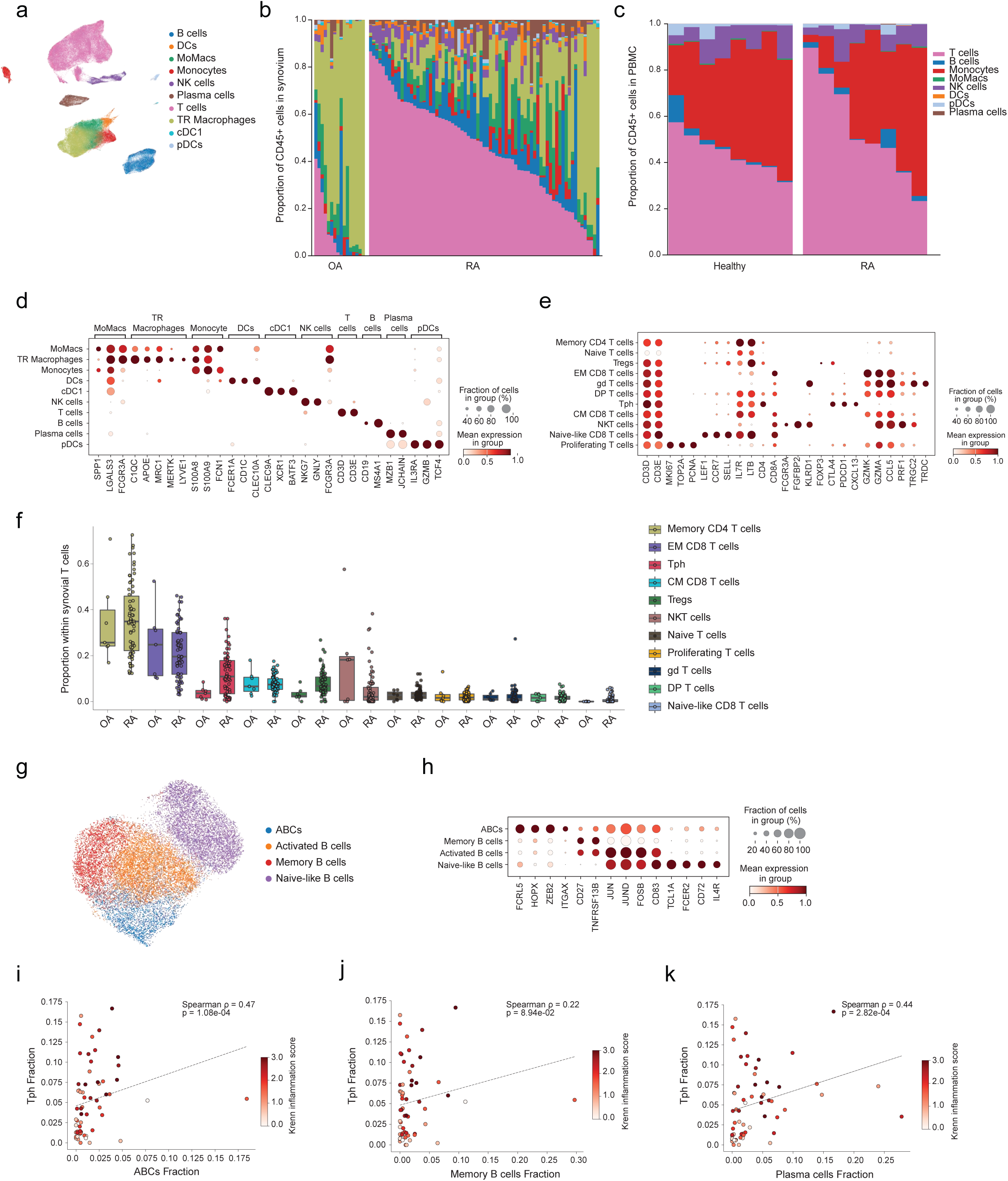
Overview of immune and disease-associated T cells in synovial tissue of RA patients. (a) Integrated immune cells UMAP, with graph-based clustering of 222,585 immune cells from the synovium and PBMC of 125 patients (91 RA, 16 OA, 18 healthy). (b) Bar graphs depicting the percentages of synovial immune subsets per individual grouped by disease. Patients with less than 100 immune cells were excluded. (c) Bar graphs depicting the percentages of PBMC immune subsets per individual grouped by disease. Patients with less than 100 immune cells were excluded. (d) Dot plot of annotation marker genes across immune clusters. Dot size indicates the fraction of cells in each cluster expressing the gene and color indicates mean expression. (e) Dot plot of annotation marker genes across T cell clusters in the dataset shown in Fig. 1b. Dot size indicates the fraction of cells in each cluster expressing the gene, and color indicates mean expression. (f) Boxplot showing the percentage of T cell subpopulation out of total synovial T cells across diseases. Each point represents a patient. (g) Integrated B cells UMAP, with graph-based clustering of 21,674 B cells from the synovium and PBMC of 107 patients (82 RA, 11 OA, 14 healthy). (h) Dot plot of annotation marker genes across B clusters. Dot size indicates the fraction of cells in each cluster expressing the gene and color indicates mean expression. (i) Spearman’s correlation testing between Tph and Age-associated B cells (ABCs) percentage of immune cells in RA patients. Each point represents a patient. Color indicates Krenn inflammation score. Spearman’s ρ = 0.47, P <0.0001. (j) Spearman’s correlation testing between Tph and memory B cells percentage of immune cells in RA patients. Each point represents a patient. Color indicates Krenn inflammation score. Spearman’s ρ = 0.22. (k) Spearman’s correlation testing between Tph and Plasma cells percentage of immune cells in RA patients. Each point represents a patient. Color indicates Krenn inflammation score. Spearman’s ρ = 0.44. P < 0.001.\

**Figure S2.**
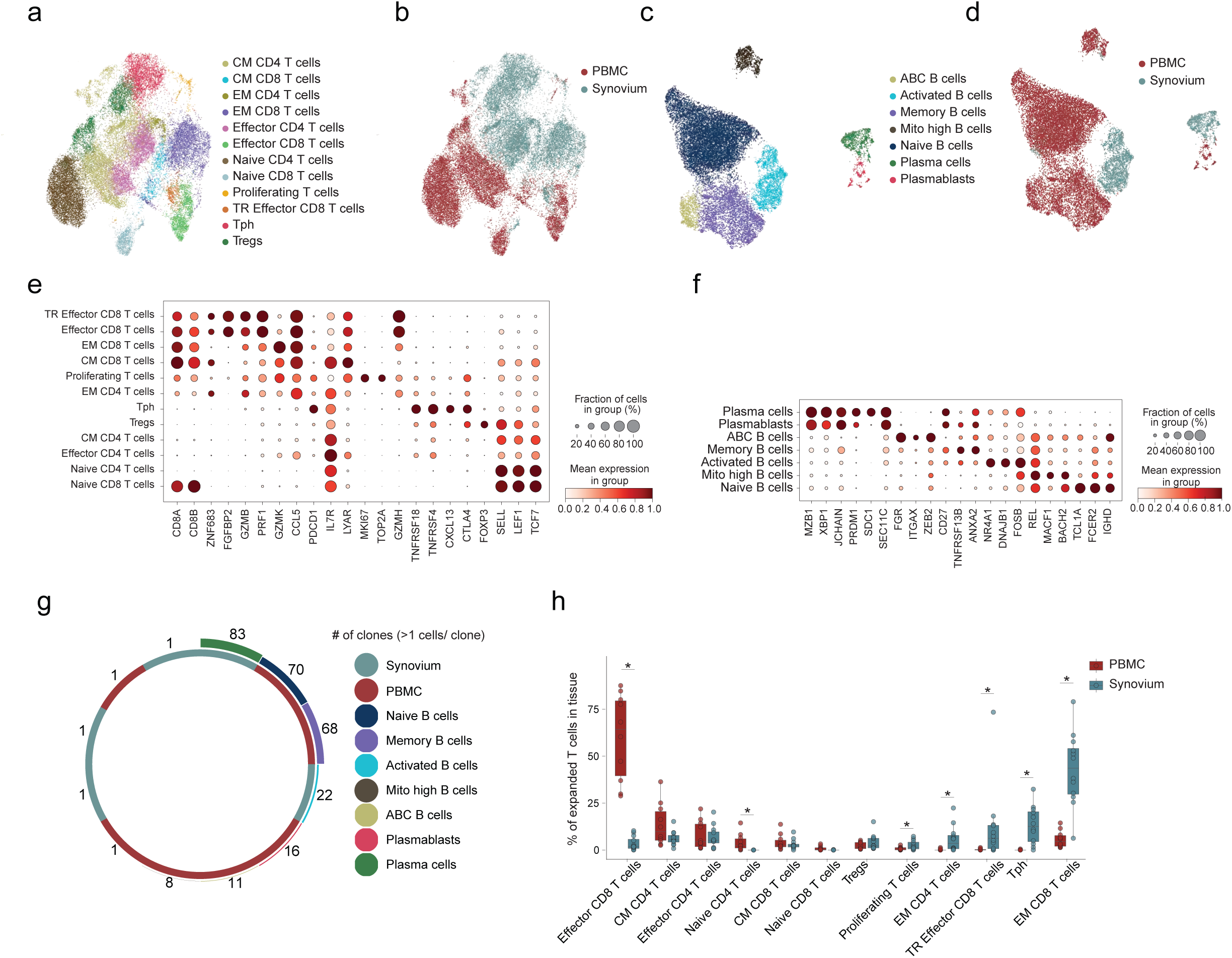
TCR-and BCR-seq of RA synovial tissue reveals disease-associated T and B Cell populations. (a) Integrated UMAP of T cells from the sc TCR seq, with graph-based clustering of 35,308 T cells from 12 synovial samples and 10 PBMC of RA patients. Right-Clone size assigned per cell. (b) Integrated UMAP of T cells from the sc TCR seq, with graph-based clustering of 35,308 T cells from 12 synovial samples and 10 PBMC of RA patients with tissue annotations. (c) Integrated UMAP of B cells from the sc BCR seq, with graph-based clustering of 21,610 B cells from the synovium and PBMC of 12 RA patients. (d) Integrated UMAP of B cells from the sc BCR seq, with graph-based clustering of 21,610 B cells from the synovium and PBMC of 12 RA patients with tissue annotations. (e) Dot plot of annotation marker genes across T cells clusters in the sc TCR dataset. Dot size indicates the fraction of cells in each cluster expressing the gene, and color indicates mean expression. (f) Dot plot of annotation marker genes across B cells clusters in the sc BCR seq dataset. Dot size indicates the fraction of cells in each cluster expressing the gene, and color indicates mean expression. (g) Circular bar plot of the number of cells with BCR clone size higher than 1, per B subpopulation, in 12 synovial samples and 10 PBMC of RA patients. (h) Boxplot showing the percentage of expanded T cell subpopulation from total expanded T cells per tissue. Each point represents a patient. (* P < 0.05).

**Figure S3.**
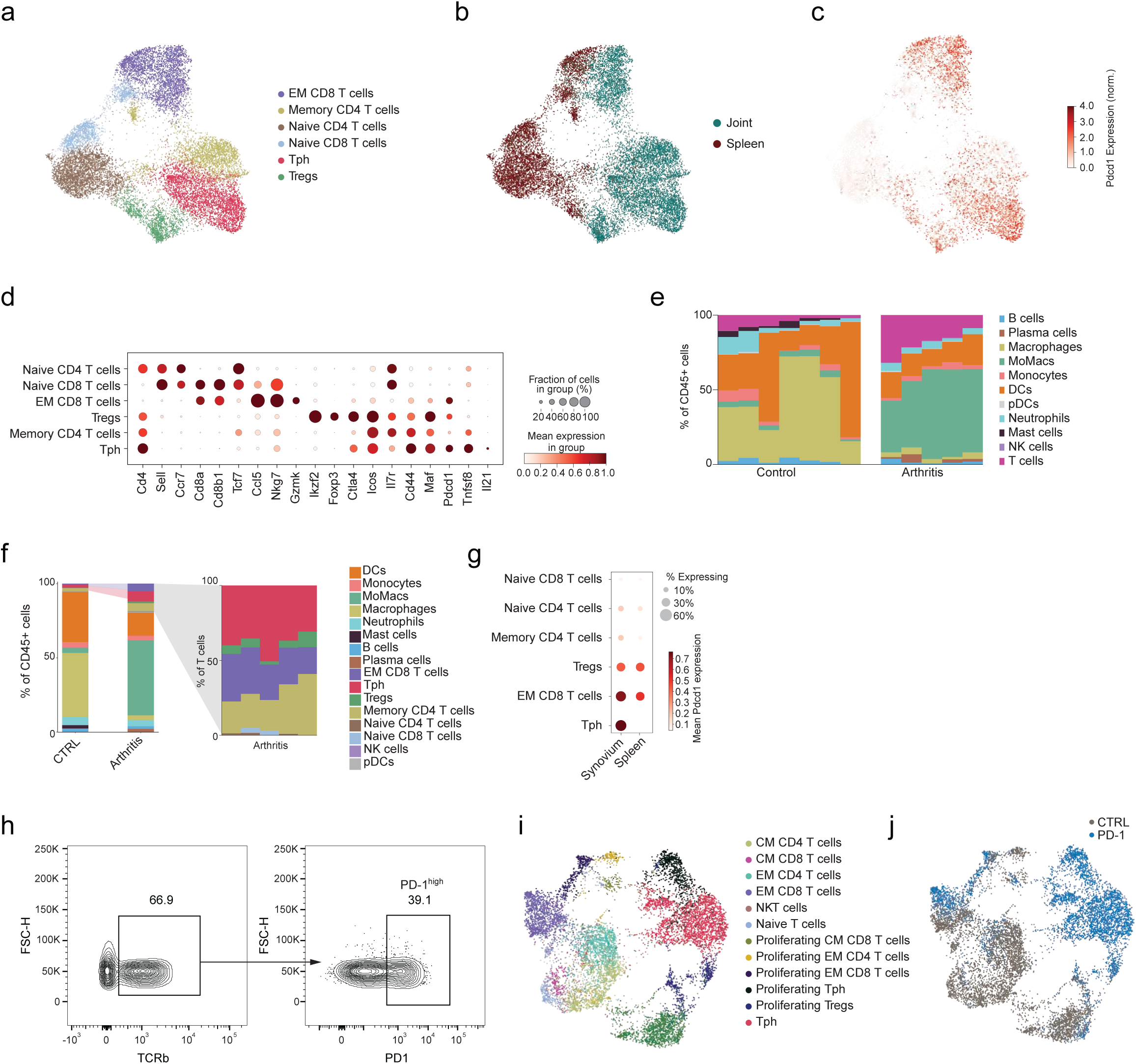
Overview of immune and disease-associated T cells in murine synovial tissue. (a) Integrated T cells UMAP from arthritis mice and controls, with graph-based clustering of 15,965 T cells from the synovium and spleen of 15 mice. (b) Integrated T cells UMAP from arthritis mice, with tissue annotations. (c) Pdcd1 normalized expression projection on integrated UMAP. (d) Dot plot of annotation marker genes across T cells. Dot size indicates the fraction of cells in each cluster expressing the gene, and color indicates mean expression. (e) Bar graphs depicting the percentages of synovial immune subsets per mouse grouped by disease state. (f) Bar graphs depicting the average percentages of synovial immune subsets per state and bar graphs depicting the percentages of T cells subsets out of total T cells per arthritis mouse. (g) Dot plot of Pdcd1 average normalized expression in the synovium and spleen of arthritis mice across T cells subsets. Dot size indicates the fraction of cells in each cluster expressing the gene, and color indicates mean expression. (h) Integrated T cells UMAP from sc TCR seq of synovium from arthritis mice, with graph-based clustering of 9,939 T cells from 11 arthritis mice. (i) Integrated T cells UMAP from sc TCR seq of synovium from arthritis mice with sample annotations.

**Figure S4.**
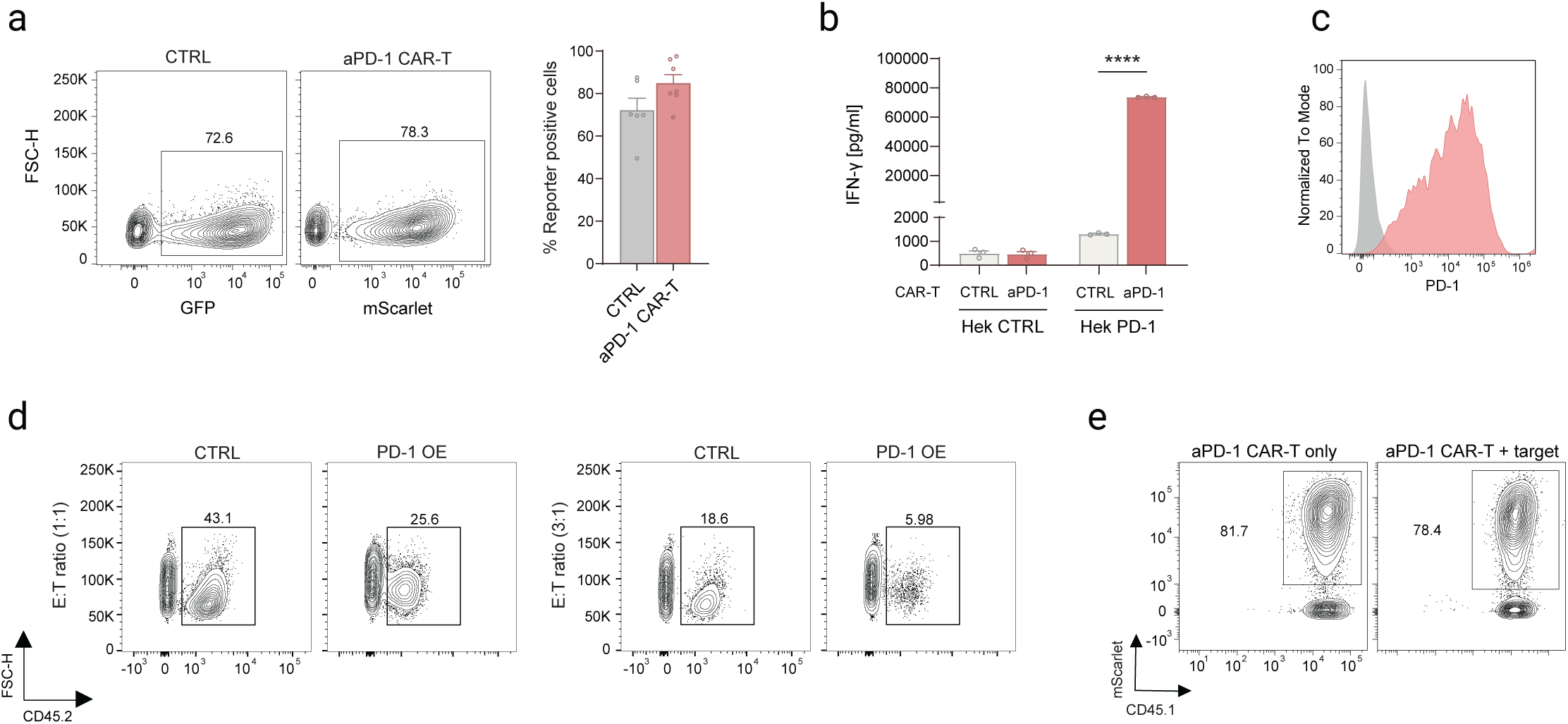
aPD-1 CAR-T cells mediate potent antigen-specific activation and selective killing of PD-1^+^ cells. (a) Representative FACS analysis of cells transduced with control GFP vector (CTRL), or aPD-1 CAR (left). Bar graphs summarize transduction efficiency (%) across experiments. (b) Mouse IFN-γ ELISA assay following 24h co-cultured of aPD-1 CAR-T and CTRL transduced T cells with Hek CTRL or Hek-PD-1 at E: T ratio of 5:1. Data are presented as mean ± SD (n = 3). Statistical significance was assessed by Two-way ANOVA (**** p < 0.0001). (c) Representative staining of T cells transduced with either a plasmid for *Pdcd1* overexpression or an empty vector. (d) Representative flow cytometry analysis of bulk T transduced with either PD-1 or empty vector cell percentages following 24h co-cultured with mouse aPD-1 CAR-T and CTRL transduced T cells at E: T ratio of 1:1 and 3:1.(e) Representative FACS staining showing aPD-1-CAR-T cell percentages (CD45.1⁺ mScarlet⁺) before and after incubation with exhausted T cells at an E:T ratio of 3:1.

**Figure S5.**
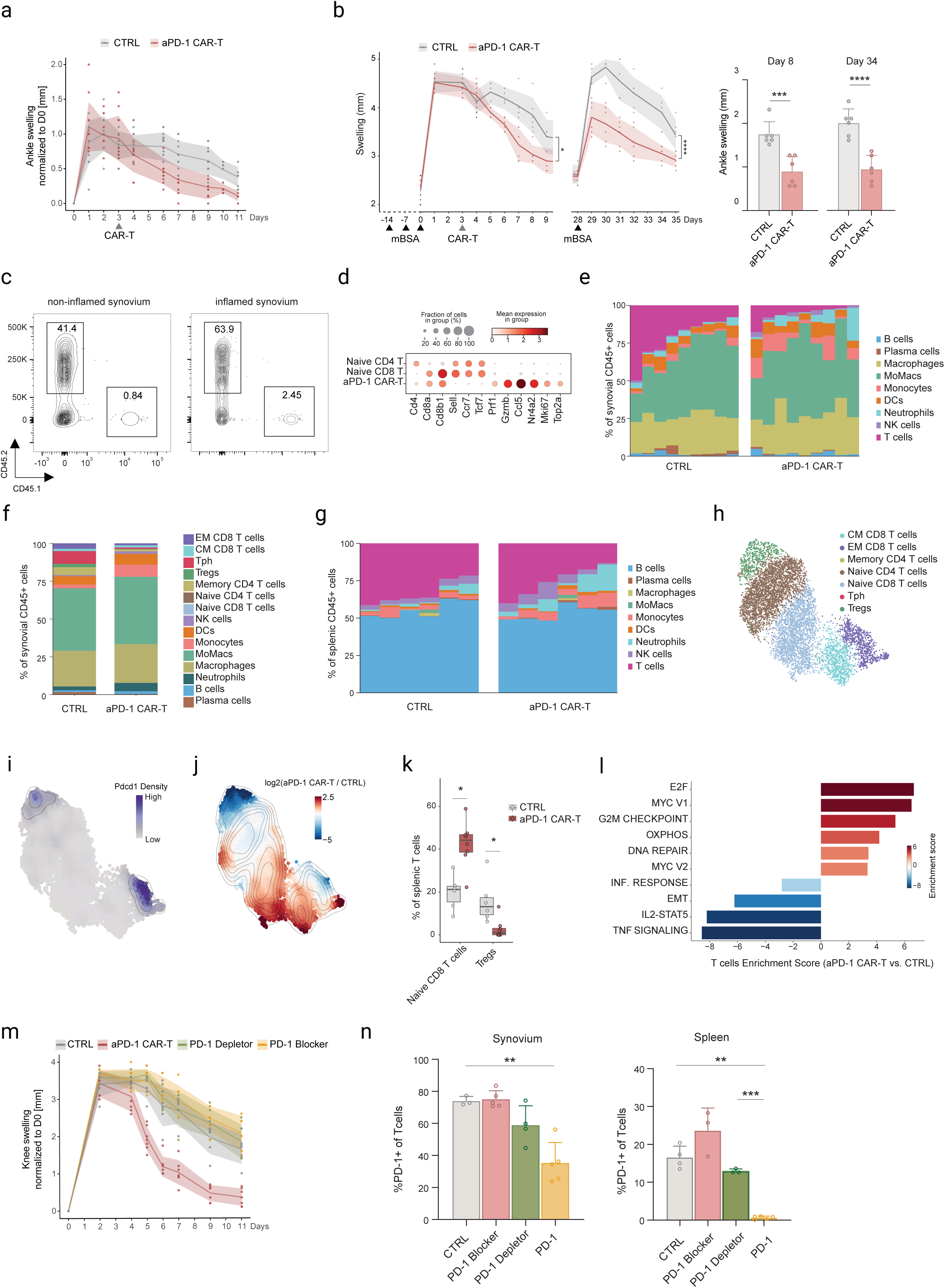
aPD-1 CAR-T mechanism of action. (a) *In vivo* efficacy of PD-1 CAR-T cell therapy in mice following arthritis induction during re-challenge phase using the mBSA arthritis model. Arthritis severity was monitored by caliper measurement of ankle joint swelling (y-axis, mm) over time following arthritis induction (x-axis, days). CAR-T cells were administered on day 3, after disease establishment. Data are presented for two groups: control-transduced T cells (CTRL), and aPD-1 CAR-T cells. n=12-13 mice/group. (b) left. *In vivo* efficacy of aPD-1 CAR-T cell therapies in mice following arthritis induction using the mBSA arthritis model. Arthritis severity was monitored by caliper measurement of joint swelling (y-axis, mm) over time following arthritis induction (x-axis, days). Disease course showed two characteristic phases: an initial inflammatory peak followed by re-challenge phase. CAR-T cells were administered on day 3, after disease establishment. Data are presented for two groups: control-transduced T cells (CTRL), and aPD-1 CAR-T cells. n=6 mice/group. Right. Summary of joint swelling during initial arthritis and re-challenge phase. Bar graphs depict mean joint swelling (mm above baseline diameter) in the two experimental groups (control, and aPD-1 CAR-T cells) at day 8 (of the initial inflammatory episode) and day 34 (re-challenge phase). One-way ANOVA (***p < 0.001). (**** p < 0.0001). (c) Representative flow cytometry analysis of CD45.1^+^ aPD-1 CAR-T cell percentages in synovial tissue of mBSA-induced arthritic joints (right) or PBS-injected normal non-inflamed synovium (left) at day 8 post-CAR injection. (d) Dot plot of selected marker genes across naïve T cells clusters and aPD-1 CAR-T cells from the synovium of arthritis mice. Dot size indicates the fraction of cells in each cluster expressing the gene, and color indicates mean expression. (e) Bar graphs depicting the percentages of synovial immune subsets per mouse grouped by condition. (f) Bar graphs depicting the average percentages of synovial immune subsets per condition with T subsets annotations. (g) Bar graphs depicting the percentages of splenic immune subsets per mouse grouped by condition. (h) Integrated T cells UMAP, with graph-based clustering of 7,190 T cells from the spleen of 18 arthritis mice, down sampled for an equal number of cells per gate in each condition. (i) Density plot showing distribution of Pdcd1 normalized expression on the splenic T cells embedding, down sampled for an equal number of cells per gate in each condition. (j) Density plots showing distribution of splenic T cells embedding per condition, down sampled for an equal number of cells per gate in each condition. (k) Boxplot showing the percentage of significantly different T subsets from total splenic T cells of arthritis mice out of CD45^+^ gate under aPD-1 CAR-T treatment compared to CTRL transduced T cells therapy. Each point represents a mouse. *P < 0.05. (l) Bar graphs of enrichment normalized scores of significant differential pathways in T cells under aPD-1 CAR-T treatment compared to CTRL transduced T cells therapy. (m) In vivo efficacy of aPD-1 CAR-T cell therapy compared with aPD-1 depleting or blocking antibodies in mice following arthritis induction during a re-challenge phase using the mBSA arthritis model. Arthritis severity was monitored by caliper measurement of knee joint swelling (y-axis, mm) over time after arthritis induction (x-axis, days). CAR-T cells or antibodies were administered on day 3, after disease establishment. Data is shown for four groups: control-transduced T cells (CTRL), aPD-1 CAR-T, anti- aPD-1 depleting or blocking antibodies. n = 6 mice per group. (n) Bar graphs depicting the frequencies of PD-1^+^ T cell in inflamed synovium (left) or spleen (right) following aPD-1 CAR-T, aPD-1 depleting or blocking antibodies, or control-transduced T cell therapies (CTRL), shown per individual mouse. (** P <0.01), (*** P < 0.001).

**Figure S6.**
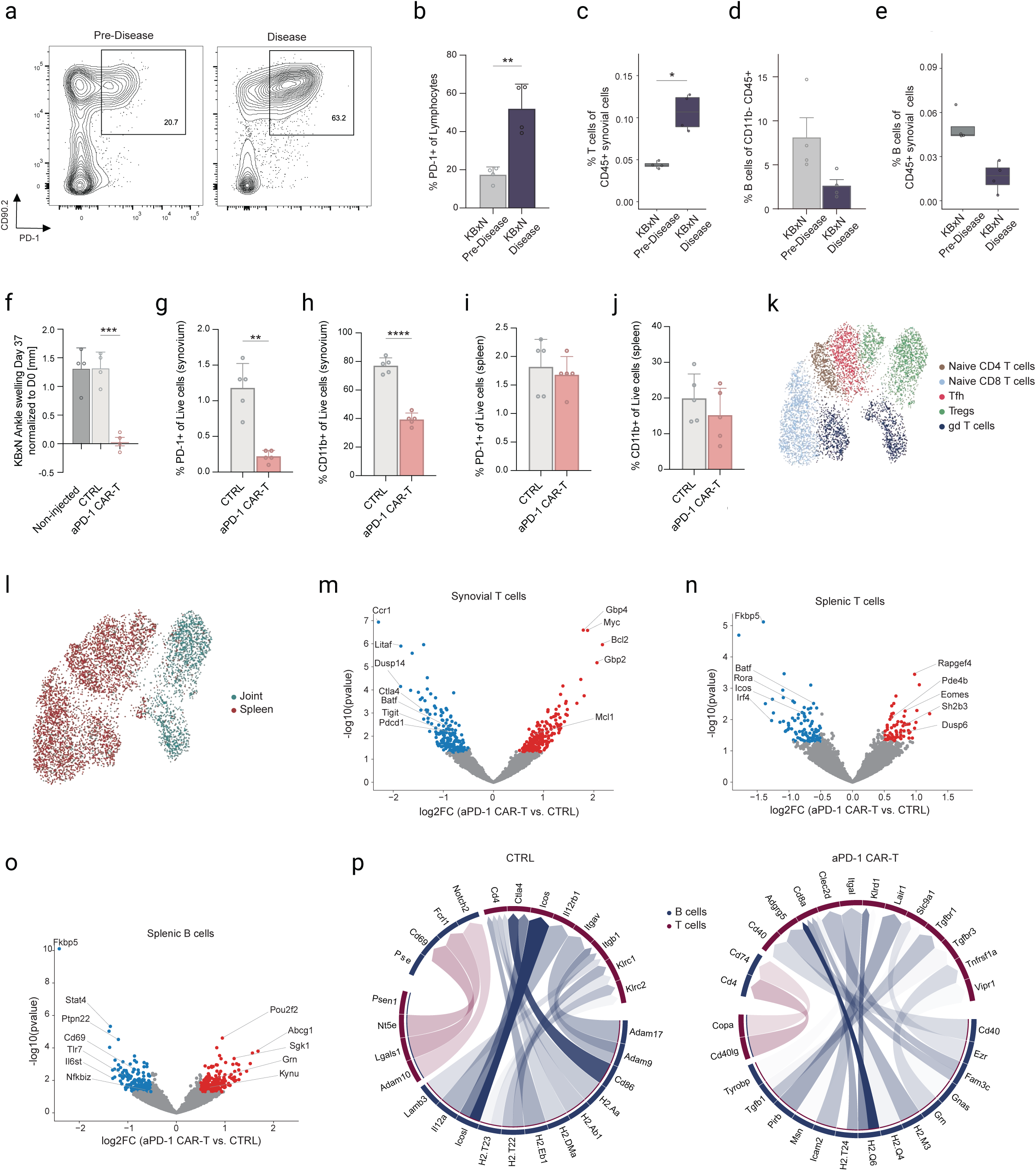
aPD-1 CAR-T cells selectively deplete PD-1^+^ disease-associated T cells in vivo, and significantly mitigate arthritis in K/BxN murine model. (a) Representative flow cytometry analysis of PD-1^+^ T cells in synovial tissue of K/BxN mice at the pre-disease stage (3 weeks of age) or after arthritis development (5 weeks). (b) Boxplot showing the percentage of PD-1^+^ T cells among total synovial lymphocytes of K/BxN mice across timepoints. Each point represents an individual mouse. (** P <0.01) (c) Boxplot showing the percentage of T cells among total synovial lymphocytes of K/BxN mice across timepoints. Each point represents an individual mouse. (*P <0.05) (d) Boxplot showing the percentage of B cells among CD11^-^CD45^+^ synovial immune cells of K/BxN mice across timepoints. Each point represents an individual mouse. (e) Boxplot showing the percentage of B cells among total synovial immune cells of K/BxN mice across timepoints. Each point represents an individual mouse. (f) Box plot showing the average ankle swelling (mm) in the K/BxN model (mean of two ankles per mouse), normalized to day 0 (pre-disease), 14 days after injection of CAR-T cells, control-transduced T cells (CTRL), or no treatment. Statistical significance was determined by one-way ANOVA (***P < 0.001). n = 4 mice per group. (g) Bar graphs depicting the percentages of PD-1^+^ T cells among live cells in K/BxN synovial tissue following aPD-1 CAR-T or control-transduced T cell (CTRL) therapies. Each dot represents an individual mouse. (**P<0.01) (h) Bar graphs show the percentage of CD11b⁺ myeloid cells among live synovial cells in K/BxN mice following aPD-1 CAR-T or CTRL-transduced T cell therapies. (****P<0.0001) (i) Bar graphs depicting the percentages of PD-1^+^ T cells among live cells in K/BxN splenic tissue following aPD-1 CAR-T or control-transduced T cell (CTRL) therapies. Each dot represents an individual mouse. (j) Bar graphs showing the percentage of CD11b⁺ myeloid cells among live splenic cells in K/BxN mice following aPD-1 CAR-T or control-transduced T cell (CTRL) therapies. (k) Integrated T cells UMAP, with graph-based clustering of 6,222 T cells of K/BxN mice. (l) Integrated T cells UMAP from K/BxN mice, with tissue annotations. (m) Volcano plot of DEGs in synovial T cells under aPD-1 CAR-T compared to CTRL treatment in K/BxN mice. Genes with adjusted P < 0.05 and |log₂FC| > 0.5 were highlighted. (n) Volcano plot of DEGs in splenic T cells under aPD-1 CAR-T compared to CTRL treatment in K/BxN mice. Genes with adjusted P < 0.05 and |log₂FC| > 0.5 were highlighted. (o) Volcano plot of DEGs in splenic B cells under aPD-1 CAR-T compared to CTRL treatment in K/BxN mice. Genes with adjusted P < 0.05 and |log₂FC| > 0.5 were highlighted. (p) Chord diagrams of ligand-receptor interaction network between splenic B cells and T cells in K/BxN mice, across treatments.

**Figure S7.**
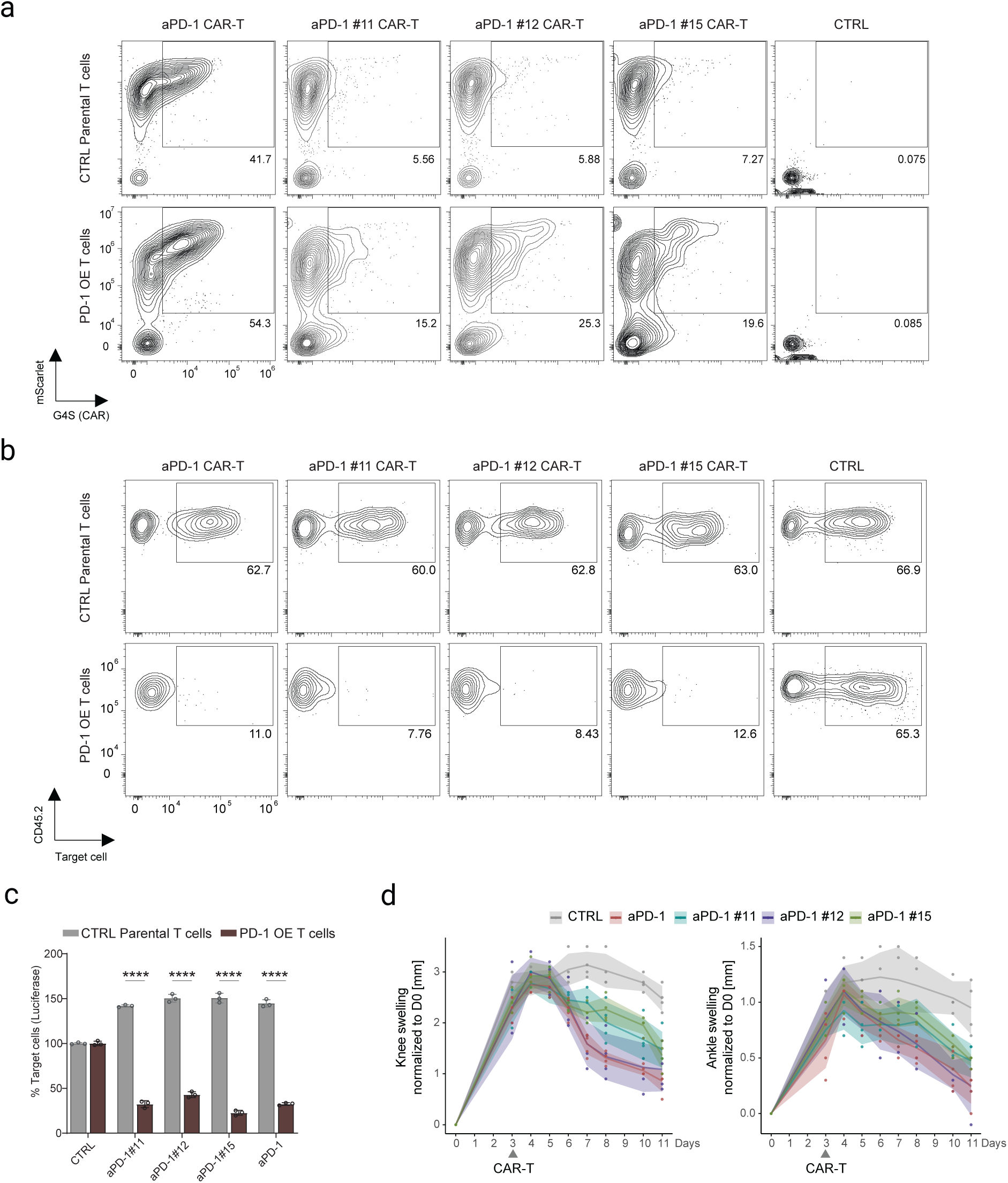
Synovium-specific regulatory elements in aPD-1 CAR-T, mechanisms of action. (a) Representative flow cytometry analysis of G4S and mScarlet expression in constitutive CD45.1^+^ aPD-1 CAR-T cells versus CD45.1^+^ regulated aPD-1#12// aPD-1 #11/ aPD-1 #15 CAR-T cells or controlled-transduced T cells (CTRL) following 24 h of incubation with CD45.2^+^ PD-1 overexpressing (OE) bulk T cells or CD45.2^+^ control parental T cells transduced with an empty vector at E: T ratio of 3:1. (b) Representative flow cytometry analysis of residual CD45.2^+^ PD-1 overexpressing (OE) bulk T cells or CD45.2^+^ control parental T cells transduced with an empty vector, following 24 h of incubation with constitutive CD45.1^+^ aPD-1 CAR-T cells or CD45.1 regulated aPD-1 #12/ aPD-1 #11/ aPD-1 #15 CAR-T cells or controlled-transduced T cells (CTRL) at an E:T ratio of 3:1. (c) Luciferase-based cytotoxicity assay of HEK293T cells expressing luciferase alone or co-expressing PD-1 and luciferase. Target cells were incubated with constitutive CAR-T cells, regulated CAR-T cells#11, #12 and #15, or control-transduced T cells (CTRL) for 24 h at an effector-to-target (E:T) ratio of 3:1. Cytotoxicity was assessed by measuring luciferase activity, with residual luciferase activity serving as a readout of target cell viability. Experiments were performed in triplicate, and data are presented as mean ± SD. ****P<0.0001 (d) *In vivo* efficacy of PD-1 CAR-T, regulated aPD-1 #11, #12, and #15 CAR, or controlled-transduced T cells (CTRL) therapies in mice following arthritis induction using the mBSA arthritis model. Arthritis severity was monitored by caliper measurement of knee (left) and ankle (right) joint swelling (y-axis, mm) over time following arthritis induction (x-axis, days). CAR-T cells or CTRL were administered on day 3, after disease establishment. Data is presented for five groups: control-transduced T cells (CTRL), aPD-1 CAR-T cells, or aPD-1#11, #12, and #15 CAR-T cells. n=4-5 mice/group.

**Figure S8.**
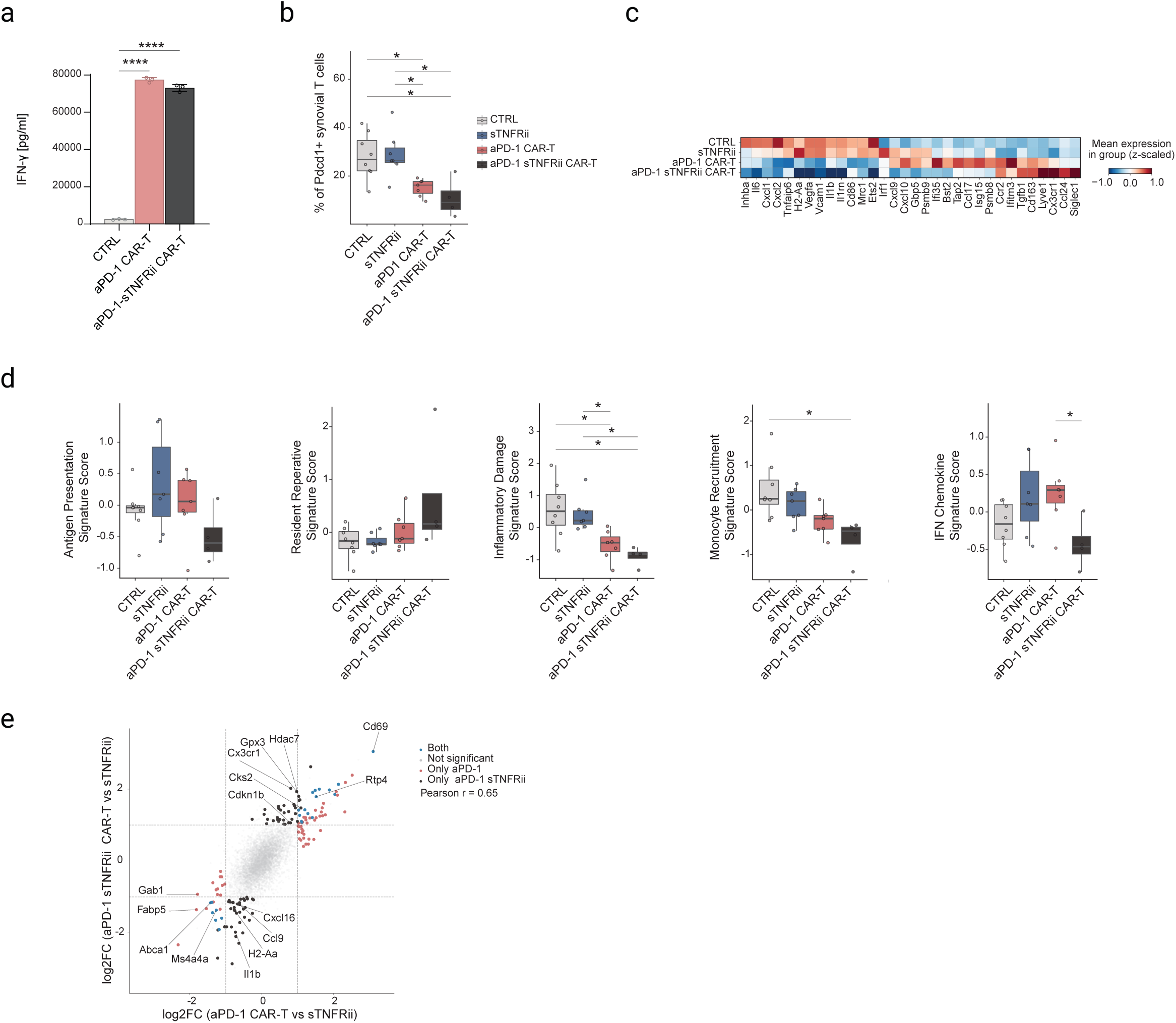
sTNFRii-secreting CAR-T cells modulate the synovial microenvironment. (a) Mouse IFN-γ ELISA assay following 24h co-cultured of aPD-1 CAR-T, aPD-1 sTNFRii CAR-T cells, and control-transduced T cells (CTRL) with Hek PD-1 cells at E:T ratio of 5:1. Data are presented as mean ± SD (n = 3). One-way ANOVA (****P<0.0001). (b) Boxplot showing percentages of PD-1^+^ synovial T cells across treatments - CTRL (n=8), sTNFRii (n=7), aPD-1 CAR-T (n=7), aPD-1 sTNFRii CAR-T (n=4). (* P < 0.05). (c) Heatmap of selected DEGs in myeloid cells across different treatments. Values are the mean normalized expression z scaled. (d) Boxplots showing different modules scores defined by genes from fig S8e across treatments- CTRL (n=8), sTNFRii (n=7), aPD-1 CAR-T (n=7), aPD-1 sTNFRii CAR-T (n=4). (* P < 0.05). (e) Scatter plot showing the log₂FC of all genes in the myeloid cells in aPD-1 CAR-T, aPD-1 sTNFRii CAR-T treated mice compared to sTNFRii treated mice. Highlighted genes have adjusted P < 0.05 and |log₂FC| > 1.

## References

1. Song, Y., Li, J. & Wu, Y. Evolving understanding of autoimmune mechanisms and new therapeutic strategies of autoimmune disorders. Signal Transduction and Targeted Therapy 2024 9:1 9, 1–40 (2024).

2. Cho, J. H. & Feldman, M. Heterogeneity of autoimmune diseases: Pathophysiologic insights from genetics and implications for new therapies. Nat. Med. 21, 730–738 (2015).

3. Pisetsky, D. S. Pathogenesis of autoimmune disease. Nat. Rev. Nephrol. 19, 1 (2023).

4. Balogh, L., Oláh, K., Sánta, S., Majerhoffer, N. & Németh, T. Novel and potential future therapeutic options in systemic autoimmune diseases. Front. Immunol. 15, 1249500 (2024).

5. Tony, H. P. et al. Safety and clinical outcomes of rituximab therapy in patients with different autoimmune diseases: Experience from a national registry (GRAID). Arthritis Res. Ther. 13, (2011).

6. Greco, R., et al. Innovative cellular therapies for autoimmune diseases: expert-based position statement and clinical practice recommendations from the EBMT practice harmonization and guidelines committee. EClinicalMedicine 69, (2024).

7. Holzer, M. T. et al. Daratumumab for autoimmune diseases: A systematic review. RMD Open 9, (2023).

8. Müller, F. et al. CD19 CAR-T cells for treatment-refractory autoimmune diseases: the phase 1/2 CASTLE basket trial. Nature Medicine 2026 32:3 32, 1142–1151 (2026).

9. Schett, G., Mackensen, A. & Mougiakakos, D. CAR T-cell therapy in autoimmune diseases. The Lancet 402, 2034–2044 (2023).

10. Schett, G. et al. Advancements and challenges in CAR T cell therapy in autoimmune diseases. Nat. Rev. Rheumatol. 20, 531–544 (2024).

11. Mackensen, A. et al. Anti-CD19 CAR T cell therapy for refractory systemic lupus erythematosus. Nat. Med. 28, 2124–2132 (2022).

12. Müller, F. et al. CD19 CAR T-Cell Therapy in Autoimmune Disease — A Case Series with Follow-up. New England Journal of Medicine 390, 687–700 (2024).

13. Rampotas, A., Richter, J., Isenberg, D. & Roddie, C. CAR-T cell therapy embarks on autoimmune disease. Bone Marrow Transplant. 60, 6–9 (2025).

14. Papatriantafyllou, M. Off-the-shelf CAR T cells for refractory autoimmunity. Nat. Rev. Rheumatol. 20, 527 (2024).

15. Scherlinger, M. et al. CAR T-cell therapy in autoimmune diseases: where are we and where are we going? Lancet Rheumatol. 7, e434–e447 (2025).

16. Mougiakakos, D. Allogeneic CAR T cells for autoimmune diseases: a glimpse into the future. Signal Transduct. Target. Ther. 9, 1–2 (2024).

17. Yang, C. et al. Allogeneic anti-CD19 CAR-T cells induce remission in refractory systemic lupus erythematosus. Cell Res. 35, 607 (2025).

18. Haghikia, A. et al. Anti-CD19 CAR T cells for refractory myasthenia gravis. Lancet Neurol. 22, 1104–1105 (2023).

19. Alivernini, S., Firestein, G. S. & McInnes, I. B. The pathogenesis of rheumatoid arthritis. Immunity 55, 2255–2270 (2022).

20. McInnes, I. B. & Schett, G. The pathogenesis of rheumatoid arthritis. N. Engl. J. Med. 365, 2205–2219 (2011).

21. Zhang, F. et al. Deconstruction of rheumatoid arthritis synovium defines inflammatory subtypes. Nature 623, 616–624 (2023).

22. Sun, W. et al. B cells inhibit bone formation in rheumatoid arthritis by suppressing osteoblast differentiation. Nature Communications 2018 9:1 9, 5127- (2018).

23. Wu, F. et al. B Cells in Rheumatoid Arthritis : Pathogenic Mechanisms and Treatment Prospects. Front. Immunol. 12, 750753 (2021).

24. Zhang, F. et al. Defining inflammatory cell states in rheumatoid arthritis joint synovial tissues by integrating single-cell transcriptomics and mass cytometry. Nat. Immunol. 20, 928–942 (2019).

25. Smolen, J. S., et al. Rheumatoid arthritis. Nat. Rev. Dis. Primers 4, (2018).

26. Papatriantafyllou, M. Two subsets of TPH cells with distinct functions in RA. Nature Reviews Rheumatology 2025 1–1 (2025) doi:10.1038/s41584-025-01300-2.

27. Ozog, S. et al. Influence of B cell-lineage targeted CAR-T cell therapy on humoral immunity and vaccine-induced antibody response. Nature Communications 2026 https://doi.org/10.1038/s41467-026-71473-1 (2026) doi:10.1038/s41467-026-71473-1.

28. Tur, C. et al. CD19-CAR T-cell therapy induces deep tissue depletion of B cells. Ann. Rheum. Dis. 84, 106–114 (2025).

29. Müller, F. et al. BCMA CAR T cells in a patient with relapsing idiopathic inflammatory myositis after initial and repeat therapy with CD19 CAR T cells. Nat. Med. 31, 1793–1797 (2025).

30. Zhang, L. et al. BCMA-Targeted T-Cell Engager for Autoimmune Hemolytic Anemia after CD19 CAR T-Cell Therapy. New England Journal of Medicine 392, 2282–2284 (2025).

31. Hsieh, E. W. Y. et al. Key considerations for advancing chimeric antigen receptor (CAR) T-cell therapy for systemic lupus erythematosus (SLE): a multi-partner/disciplinary working group perspective. RMD Open 11, 5866 (2025).

32. Freeley, M. CAR T Cell Therapy for Rheumatoid Arthritis. Clin. Rev. Allergy Immunol. 68, (2025).

33. Lidar, M. et al. CD-19 CAR-T cells for polyrefractory rheumatoid arthritis. Ann. Rheum. Dis. 84, 370–372 (2025).

34. Li, Y. et al. Fourth-generation chimeric antigen receptor T-cell therapy is tolerable and efficacious in treatment-resistant rheumatoid arthritis. Cell Research 2025 35:3 35, 220–223 (2025).

35. Xavier, R. J. & Rioux, J. D. Genome-wide association studies: A new window into immune-mediated diseases. Nat. Rev. Immunol. 8, 631–643 (2008).

36. Okada, Y. et al. Genetics of rheumatoid arthritis contributes to biology and drug discovery. Nature 2013 506:7488 506, 376–381 (2013).

37. Dunlap, G. et al. Clonal associations between lymphocyte subsets and functional states in rheumatoid arthritis synovium. Nature Communications 2024 15:1 15, 4991- (2024).

38. Argyriou, A. et al. Single cell sequencing identifies clonally expanded synovial CD4+ TPH cells expressing GPR56 in rheumatoid arthritis. Nat. Commun. 13, (2022).

39. Binvignat, M., et al. Single-cell RNA-Seq analysis reveals cell subsets and gene signatures associated with rheumatoid arthritis disease activity. JCI Insight 9, (2024).

40. Masuo, Y., et al. Stem-like and effector peripheral helper T cells comprise distinct subsets in rheumatoid arthritis. Sci. Immunol. 10, eadt3955 (2025).

41. Rao, D. A. et al. Pathologically expanded peripheral T helper cell subset drives B cells in rheumatoid arthritis. Nature 542, 110–114 (2017).

42. Weinand, K. et al. The chromatin landscape of pathogenic transcriptional cell states in rheumatoid arthritis. Nature Communications 2024 15:1 15, 4650- (2024).

43. Jonsson, A. H. et al. Granzyme K+ CD8 T cells form a core population in inflamed human tissue. Sci. Transl. Med. 14, eabo0686 (2022).

44. Philpott, H. T. et al. Association of Synovial Innate Immune Exhaustion With Worse Pain in Knee Osteoarthritis. Arthritis and Rheumatology 77, 664–676 (2025).

45. Lopez, R. et al. DestVI identifies continuums of cell types in spatial transcriptomics data. Nat. Biotechnol. 40, 1360–1369 (2022).

46. Krenn, V. et al. Synovitis score: Discrimination between chronic low-grade and high-grade synovitis. Histopathology 49, 358–364 (2006).

47. Frey, O. et al. The role of regulatory T cells in antigen-induced arthritis: aggravation of arthritis after depletion and amelioration after transfer of CD4+CD25+ T cells. Arthritis Res. Ther. 7, R291 (2005).

48. van den Berg, W. B., Joosten, L. A. B. & van Lent, P. L. E. M. Murine antigen-induced arthritis. Methods Mol. Med. 136, 243–253 (2007).

49. Jaitin, D. A. et al. Massively parallel single-cell RNA-seq for marker-free decomposition of tissues into cell types. Science (1979). 343, 776–779 (2014).

50. Yofe, I. et al. Spatial and Temporal Mapping of Breast Cancer Lung Metastases Identify TREM2 Macrophages as Regulators of the Metastatic Boundary. Cancer Discov. 13, 2610–2631 (2023).

51. Dahan, R. et al. FcγRs Modulate the Anti-tumor Activity of Antibodies Targeting the PD-1/PD-L1 Axis. Cancer Cell 28, 285–295 (2015).

52. Yamazaki, T. et al. Blockade of B7-H1 on Macrophages Suppresses CD4+ T Cell Proliferation by Augmenting IFN-γ-Induced Nitric Oxide Production. The Journal of Immunology 175, 1586–1592 (2005).

53. Monach, P. A., Mathis, D. & Benoist, C. The K/BxN arthritis model. Curr. Protoc. Immunol. **Chapter** 15, (2008).

54. Ditzel, H. J. The K/BxN mouse: a model of human inflammatory arthritis. Trends Mol. Med. 10, 40–45 (2004).

55. Tafuri, A. et al. ICOS is essential for effective T-helper-cell responses. Nature 409, 105–109 (2001).

56. Chen, J. et al. NR4A transcription factors limit CAR T cell function in solid tumours. Nature 567, 530–534 (2019).

57. Gallagher, M. P. et al. Hierarchy of signaling thresholds downstream of the T cell receptor and the Tec kinase ITK. Proc. Natl. Acad. Sci. U. S. A. 118, e2025825118 (2021).

58. Wong, M. et al. TNFα blockade in human diseases: Mechanisms and future directions. Clin. Immunol. 126, 121 (2007).

59. Kalliolias, G. D. & Ivashkiv, L. B. TNF biology, pathogenic mechanisms and emerging therapeutic strategies. Nat. Rev. Rheumatol. 12, 49 (2015).

60. van Loo, G. & Bertrand, M. J. M. Death by TNF: a road to inflammation. Nat. Rev. Immunol. 23, 289 (2022).

61. Goldenberg, M. M. Etanercept, a novel drug for the treatment of patients with severe, active rheumatoid arthritis. Clin. Ther. 21, 75–87 (1999).

62. Giavridis, T. et al. CAR T cell-induced cytokine release syndrome is mediated by macrophages and abated by IL-1 blockade. Nat. Med. 24, 731–738 (2018).

63. Kearsley-Fleet, L. et al. Biologic refractory disease in rheumatoid arthritis: results from the British Society for Rheumatology Biologics Register for Rheumatoid Arthritis. Ann. Rheum. Dis. 77, 1405–1412 (2018).

64. David, P. et al. Poly-Refractory Rheumatoid Arthritis: An Uncommon Subset of Difficult to Treat Disease With Distinct Inflammatory and Noninflammatory Phenotypes. Arthritis Rheumatol. 76, 510–521 (2024).

65. Buch, M. H., Eyre, S. & McGonagle, D. Persistent inflammatory and non-inflammatory mechanisms in refractory rheumatoid arthritis. Nat. Rev. Rheumatol. 17, 17–33 (2021).

66. Fraietta, J. A. et al. Determinants of response and resistance to CD19 chimeric antigen receptor (CAR) T cell therapy of chronic lymphocytic leukemia. Nature Medicine 2018 24:5 24, 563–571 (2018).

67. Suga, K. et al. Single-Cell Analysis Reveals Peripheral Helper T Cells in Rheumatoid Arthritis-Related Interstitial Lung Disease. Arthritis Rheumatol. https://doi.org/10.1002/ART.70066 (2026) doi:10.1002/ART.70066.

68. Seki, N. et al. Cytotoxic Tph subset with low B-cell helper functions and its involvement in systemic lupus erythematosus. Communications Biology 2024 7:1 7, 277- (2024).

69. Chen, W., Yang, F. & Lin, J. Tph Cells Expanded in Primary Sjögren’s Syndrome. Front. Med. (Lausanne). 9, 900349 (2022).

70. Jones, G. W., Hill, D. G., Sime, K. & Williams, A. S. In Vivo Models for Inflammatory Arthritis. https://doi.org/10.1007/978-1-4939-7568-6_9 doi:10.1007/978-1-4939-7568-6_9.

71. Murphy, E. P., Dobson, A. D., Keller, C. & Conneely, O. M. Differential Regulation of Transcription by the NURR1/NUR77 Subfamily of Nuclear Transcription Factors. Gene Expr. 5, 169 (2018).

72. Grant, C. E., Bailey, T. L. & Noble, W. S. FIMO: scanning for occurrences of a given motif. Bioinformatics 27, 1017–1018 (2011).

73. Yagel, G. et al. Tumor-antigen-independent targeting of solid tumors by armored macrophage-directed anti-TREM2 CAR T cells. Cancer Cell 44, 519–533.e10 (2026).

74. Wolf, F. A., Angerer, P. & Theis, F. J. SCANPY: Large-scale single-cell gene expression data analysis. Genome Biol. 19, 1–5 (2018).

75. Love, M. I., Huber, W. & Anders, S. Moderated estimation of fold change and dispersion for RNA-seq data with DESeq2. Genome Biol. 15, 1–21 (2014).

76. Badia-I-Mompel, P. et al. decoupleR: ensemble of computational methods to infer biological activities from omics data. Bioinformatics Advances 2, (2022).

77. Wolock, S. L., Lopez, R. & Klein, A. M. Scrublet: Computational Identification of Cell Doublets in Single-Cell Transcriptomic Data. Cell Syst. 8, 281 (2019).

78. Sturml, G., et al. Scirpy: a Scanpy extension for analyzing single-cell T-cell receptor-sequencing data. Bioinformatics 36, 4817–4818 (2020).

79. Bredikhin, D., Kats, I. & Stegle, O. MUON: multimodal omics analysis framework. Genome Biol. 23, (2022).

80. Browaeys, R., Saelens, W. & Saeys, Y. NicheNet: modeling intercellular communication by linking ligands to target genes. Nat. Methods 17, 159–162 (2020).

81. Muzellec, B., Teleńczuk, M., Cabeli, V. & Andreux, M. PyDESeq2: a python package for bulk RNA-seq differential expression analysis. Bioinformatics 39, (2023).

